# Cross-species molecular mapping of the photoreceptor sensory cilium and periciliary structures identifies conserved and species-specific architectural features

**DOI:** 10.64898/2026.05.24.727487

**Authors:** Kei Takahashi, Mayuna Obayashi, Raghavi Sudharsan, William A. Beltran

**Author notes:** Corresponding Author: Kei Takahashi, Division of Experimental Retinal Therapies, Department of Clinical Sciences & Advanced Medicine, School of Veterinary Medicine, University of Pennsylvania, 3900 Delancey Street, Philadelphia, PA, 19104, USA.; William A. Beltran, Division of Experimental Retinal Therapies, Department of Clinical Sciences & Advanced Medicine, School of Veterinary Medicine, University of Pennsylvania, 3900 Delancey Street, Philadelphia, PA, 19104, USA. **Commercial Relationships Disclosures:** K.T. None; M.O. None; R.S. None; W.A.B. None.

## Abstract

Photoreceptor sensory cilia comprise a microtubule-based structural core and the phototransductive outer segment. The microtubule-based ciliary axoneme extends from the basal body through the connecting cilium into the outer segment, thereby linking the inner and outer segments. Around this central axis, specialized periciliary structures are distributed across the apical inner segment and basal outer segment. Yet how these ciliary and periciliary structures differ between rods and cones and across mammals remains incompletely defined. Using ultrastructure expansion microscopy, we compared their nanoscale molecular organization across canine, non-human primate (NHP), and human retinas. Key rod–cone differences in ciliary organization were broadly conserved among all three species, whereas the scale and morphology of individual ciliary subcompartments differed between species. Periciliary structures showed even greater species-specific diversification, particularly in rods. Human photoreceptors shared several structural features with NHP photoreceptors but also displayed inter-individual variability in accessory inner segment-like structure. These findings refine current understanding of photoreceptor sensory cilia and associated periciliary structures in large mammals, revealing that conserved rod–cone differences coexist with pronounced species-specific structural and molecular specializations.

## Introduction

Photoreceptors are light-sensing retinal neurons distinguished by a highly specialized ciliary compartment, known as the photoreceptor sensory cilium (PSC). In vertebrate retinas, photoreceptors are broadly classified into rods and cones, which are specialized for dim-light and daylight/color vision, respectively (Lamb, 2016; Molday and Moritz, 2015). In both rod and cone photoreceptors, the PSC comprises a microtubule-based structural core together with the outer segment (OS), a highly specialized membrane organelle dedicated to phototransduction. The microtubule-based structural core includes the daughter centriole (DC), basal body (BB), and ciliary axoneme, which extends from the distal end of the BB through the connecting cilium (CC) and continues into the OS (**Fig. 1A**). Immediately distal to the CC, the ciliary axoneme widens at the OS base, adjacent to the site of nascent disc or lamella formation in the proximal OS, forming a domain termed “the bulge region” before continuing as the distal axoneme (Faber et al., 2023; Mercey et al., 2022). The OS contains stacked membrane discs or lamellae that undergo continual renewal, requiring a substantial and continuous supply of membrane and proteins from the inner segment (IS) through the CC (Barnes et al., 2021). Together, the DC, BB, and ciliary axoneme form an integrated ciliary framework that links the IS to the membrane-rich OS and supports OS formation and maintenance.

**Figure 1.**
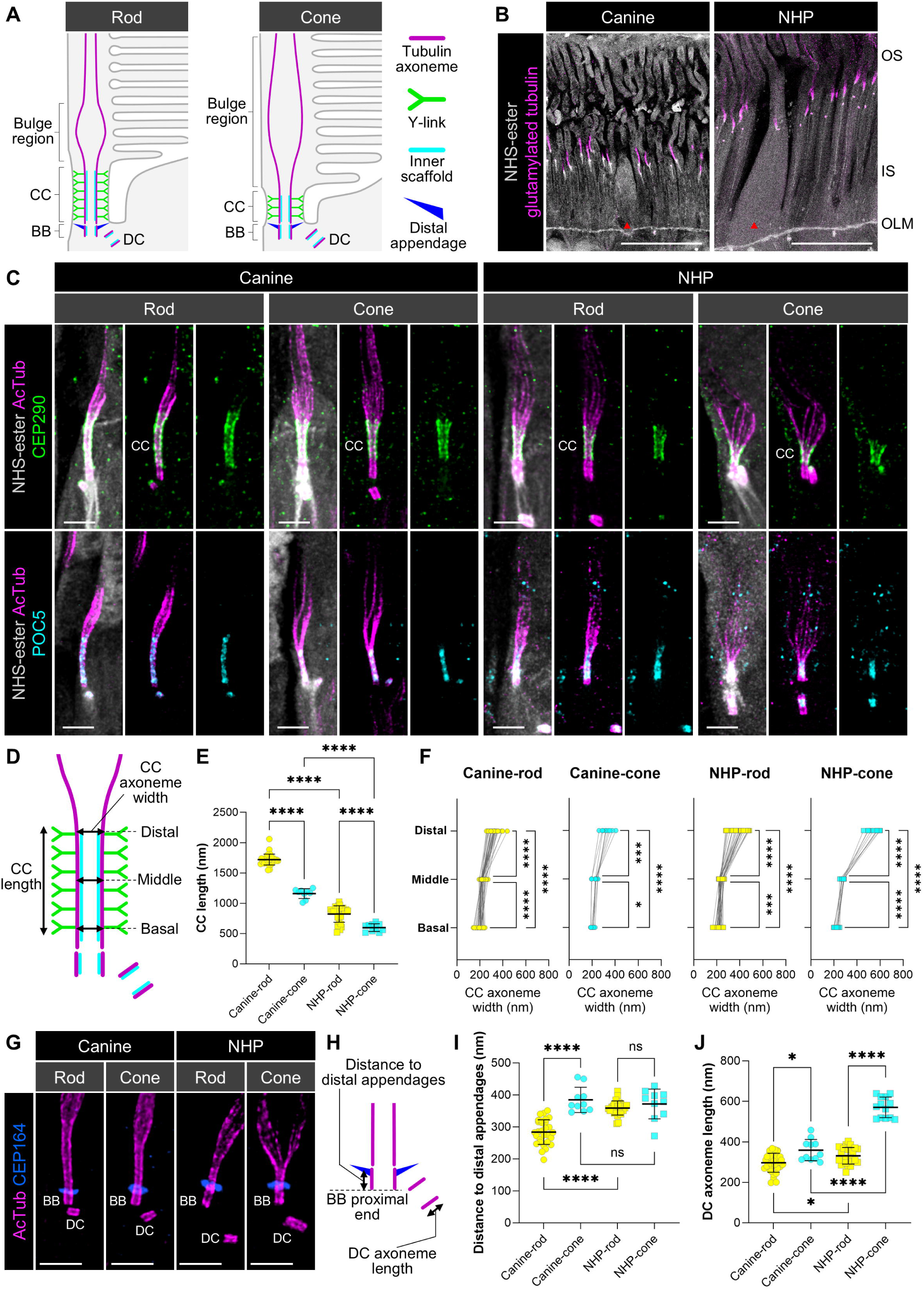
Cross-species analysis of the basal region of the PSC in canine and NHP photoreceptors. **(A)** Schematic diagram of rod (left) and cone (right) photoreceptor sensory cilia (PSCs). In this manuscript, basal body (BB) refers to the mother centriole-derived ciliary basal part, whereas daughter centriole (DC) refers to the adjacent centriole. **(B)** Low-magnification confocal images showing an overview of the outer retina in canine (left) and non-human primate (NHP; right) eyes. Photoreceptors are outlined by N-hydroxysuccinimide (NHS)-ester pan-staining (gray). Overall, NHP photoreceptors exhibit longer IS/OS profiles and greater transverse width/diameter at the IS level than canine photoreceptors. PSC axoneme is visualized by glutamylation immunolabeling (magenta). All images are shown as maximum intensity projections. Scale bars, 10 μm, corrected for the expansion factor. **(C)** Representative confocal images showing molecular localization in the connecting cilium (CC) of PSCs in rod and cone photoreceptors from canine and NHP retinas. Upper panels show CEP290 (green), a Y-link component of the CC, whereas lower panels show POC5 (cyan), a component of the tubulin inner scaffold. The PSC axoneme is visualized by acetylated α-tubulin (AcTub) immunolabeling (magenta). All images are shown as maximum intensity projections. Scale bars, 1 μm, corrected for the expansion factor. **(D)** Schematic diagram illustrating measurement of CC length and CC axoneme width. CC length was measured based on the CEP290-labeled Y-link distribution, whereas CC axoneme width was measured based on the AcTub-labeled axoneme at the corresponding region. **(E, F)** Quantification of CC length (E), measured from CEP290 immunolabeling, and CC axoneme width (F), measured from AcTub immunolabeling, in canine and NHP retinas (canine, circles; NHP, squares). Central lines indicate the mean, and error bars represent ± SD. **(G)** Representative images highlighting the basal region of PSCs in canine and NHP photoreceptors. Distal appendages are visualized by CEP164 labeling (blue). All images are shown as maximum intensity projections. Scale bars, 1 μm, corrected for the expansion factor. All microscopy images in this figure are U-ExM images. **(H)** Schematic diagram illustrating measurement of the distance from the proximal end of the BB to the distal appendages and of DC axoneme length. **(I, J)** Quantification of the distance to the distal appendages (I) and DC axoneme length (J) in canine and NHP retinas (canine, circles; NHP, squares). Central lines indicate the mean, and error bars represent ± SD. Data were obtained from three canine and two NHP eyes. \**P* < 0.05, *** *P* < 0.001, **** *P* < 0.0001, as assessed by Welch’s one-way ANOVA followed by Dunnett’s T3 multiple-comparison test.

In the periciliary region surrounding the apical IS, the PSC is accompanied by plasma membrane specializations, including a ciliary pocket and adjacent periciliary membrane domain (Sánchez-Bellver et al., 2021; Wensel et al., 2021). Although the ciliary pocket/periciliary membrane region shares conceptual similarities with membrane subdomains at the base of primary cilia (Anvarian et al., 2019; Mill et al., 2023), the elaborated photoreceptor periciliary structures, including the IS/OS junction and calyceal processes, represents a photoreceptor-specialized architecture. These periciliary domains are further distinguished by the compartmentalized distribution of Usher syndrome (USH)-associated proteins, which are disease-associated proteins but also serve as useful molecular markers of photoreceptor periciliary organization. In many vertebrate photoreceptors, USH type 2 (USH2)-associated proteins localize to the periciliary membrane region, whereas USH type 1 (USH1)-associated proteins are enriched at the IS/OS junction and/or calyceal process-associated membrane contacts (El-Amraoui and Petit, 2014; Patel et al., 2026; Sahly et al., 2012; Takahashi et al., 2025).

Accumulating evidence from studies of these specialized photoreceptor structures has begun to reveal rod-cone variation and species-dependent differences at the ultrastructural and molecular levels, raising the question of how these architectural features are conserved or diversified across mammals. Recent application of ultrastructure expansion microscopy (U-ExM) to the mature canine retina revealed striking differences between rod and cone photoreceptors across multiple PSC-associated subcompartments, including CC length, bulge region architecture, ciliary rootlet morphology and the organization of USH proteins (Takahashi et al., 2025). These findings established that rod-cone differences in ciliary molecular architecture can be resolved in formaldehyde-fixed frozen archival tissue of the adult large-animal retina at nanoscale resolution by utilizing U-ExM. However, whether such rod- and cone-specific structural features are conserved across mammals, particularly in non-human primates (NHP), which are key translational models with close relevance to human photoreceptor biology, and in human photoreceptors themselves, has not been directly and systematically examined. Although systematic cross-species comparison of the microtubule-based PSC axoneme remains limited, associated periciliary structures, including actin-rich calyceal processes, are already known to exhibit pronounced interspecies and rod-cone variation, with calyceal processes being well developed in NHP and human photoreceptors but absent or vestigial in rodents (Sahly et al., 2012; Verschueren et al., 2022). In addition, recent ultrastructural analysis identified a human rod-specific accessory inner segment, further supporting substantial specialization of structures surrounding the PSC across species and between rods and cones (Lewis et al., 2025a). Here, by applying U-ExM to formaldehyde-fixed frozen archival retinal samples from mature canine, NHP and human eyes, we compare PSC architecture and associated periciliary structures between rod and cone photoreceptors and across species, with the aim of refining current concepts of large-mammalian photoreceptor ciliary organization.

## Results

### PSC architecture shows shared rod–cone patterns and species-dependent features in canine and NHP photoreceptors

In our recent study, we showed that the microtubule-based structural core of the young adult canine PSC differs markedly between rods and cones, with cones exhibiting shorter CCs, longer BBs and DCs, and a more extended bulge region than rods (**Fig. 1A**). To enable direct structural comparison between canine and NHP PSCs, we first established a common expansion factor (EF) using retinal samples that had undergone identical fixation and cryopreservation procedures. Following previous PSC studies using U-ExM (Faber et al., 2023; Mercey et al., 2024; Mercey et al., 2022), we estimated the EF from the width of the BB axoneme, based on the assumption that mother centriole width (231,3 nm) is conserved across mammalian species. BB axoneme width was comparable across species and between rods and cones, and an average EF of 3.64 was therefore used for all subsequent measurements (**Fig. S1**).

To further minimize potential confounding factors in cross-species comparisons, we also considered whether PSC dimensions might vary according to retinal location. Because the canine values reported previously were derived from photoreceptors sampled across multiple retinal regions (Takahashi et al., 2025), we compared CC length between superior central and superior peripheral regions in the canine retina. CCs were modestly but significantly longer in both rods and cones in the superior central region than in the peripheral region (**Fig. S2**). Therefore, to maintain regional consistency and avoid potential confounding factor from macular or foveal cone specialization, we used anatomically matched superior central retinal regions outside the macular/foveal region for all cross-species analyses described below.

Using N-hydroxysuccinimide (NHS)-ester counterstaining (M’Saad and Bewersdorf, 2020), which broadly labels cellular proteins, we were able to visualize the overall photoreceptor morphology and distinguish rods from cones based on their structural features without additional rod-and cone-specific labeling. In low-magnification U-ExM images, NHP photoreceptors showed larger IS/OS profiles than canine photoreceptors, including greater longitudinal length and transverse width/diameter in the sampled retinal region (**Fig. 1B**). This difference was also observed in non-expanded retinal sections, indicating that the apparent size difference was not generated by the expansion procedure (**Fig. S3**). These observations are consistent with previous conventional histological studies showing larger primate photoreceptor IS/OS profiles compared with those of rodents and other mammals, including dogs (Booler et al., 2022; Grünert and Martin, 2020). In contrast, BB axoneme width was comparable across species and between rods and cones, consistent with the use of this structure as an internal reference for expansion factor estimation (**Fig. S1**). Closer examination of individual PSC subcompartments, however, revealed clear rod-cone variation and species-dependent differences.

We first focused on the CC, measuring its length based on centrosomal protein of 290 kDa (CEP290) signal distribution, a representative Y-link-associated marker of this compartment (**Fig. 1D**). As in canine retina, NHP cones exhibited significantly shorter CCs than NHP rods. In addition, CCs were significantly shorter in NHP than in canine photoreceptors in both rods and cones (**Fig. 1E**). These rod-cone and species-dependent differences in CC length were further supported by labeling with additional CC-associated markers, including spermatogenesis associated 7 (SPATA7) and glutamylation signals located outside the ciliary axoneme (**Fig. S4**). By contrast, the width of the axoneme at the basal CC region was similar across species and between rods and cones and was consistent with the previously reported mouse CC axoneme width of approximately 200 nm (Gilliam et al., 2012) (**Fig. 1F**). However, in both species, axoneme width was not uniform along the CC and gradually increased toward the distal end in both rods and cones. Notably, the distal widening of the CC was more prominent in NHP than in canine photoreceptors (**Fig. 1F**).

We then examined the BB and DC region in canine and NHP retinas (**Fig. 1G, H**). In canine cones, the distance from the proximal end of the BB to the centrosomal protein of 164 kDa (CEP164)-positive distal appendages was significantly greater than in canine rods, whereas no significant rod–cone difference was detected in NHP photoreceptors (**Fig. 1I**). In contrast, the feature of longer DCs in cones than in rods was shared between canine and NHP PSCs. Among all groups, NHP cones exhibited the longest DCs, approximately twice the length measured in the other groups (**Fig. 1J**). We also noted that NHP rods showed a larger apparent separation between the BB and DC than the other photoreceptor groups (**Fig. 1C, G**), although BB–DC distance was not formally quantified and was not further interpreted.

To define the structural organization of the distal CC/proximal OS axoneme region, we next labeled retinitis pigmentosa 1 protein (RP1), a photoreceptor axoneme-associated protein, and Leber congenital amaurosis 5 protein (LCA5/lebercilin), a cilium-associated protein enriched in the bulge region (Takahashi et al., 2025). These proteins served as molecular markers of the distal axoneme and bulge region, respectively (**Fig. 2A, B**). The distances from the proximal end of the BB to RP1 and LCA5 signals reflected the differences in CC length between rods and cones and across species (**Fig. 2C, D**). In addition, qualitative morphological observations of RP1 and LCA5 distribution indicated that NHP photoreceptors possess a more pronounced bulge morphology than canine photoreceptors. To examine this region in more detail, we performed axial-view imaging of the region distal to the CC end (**Fig. 2E, F**). In canine photoreceptors, the bulge morphology appeared more pronounced in cones than in rods, consistent with the quantitative findings in previous report (Takahashi et al., 2025). In NHP photoreceptors, both rods and cones displayed more pronounced bulge morphology than canine rods, with NHP cones showing the most pronounced morphology among all groups. In addition, the bulge region in canine cones and NHP photoreceptors appeared more elliptical than in canine rods (**Fig. 2F**).

**Figure 2.**
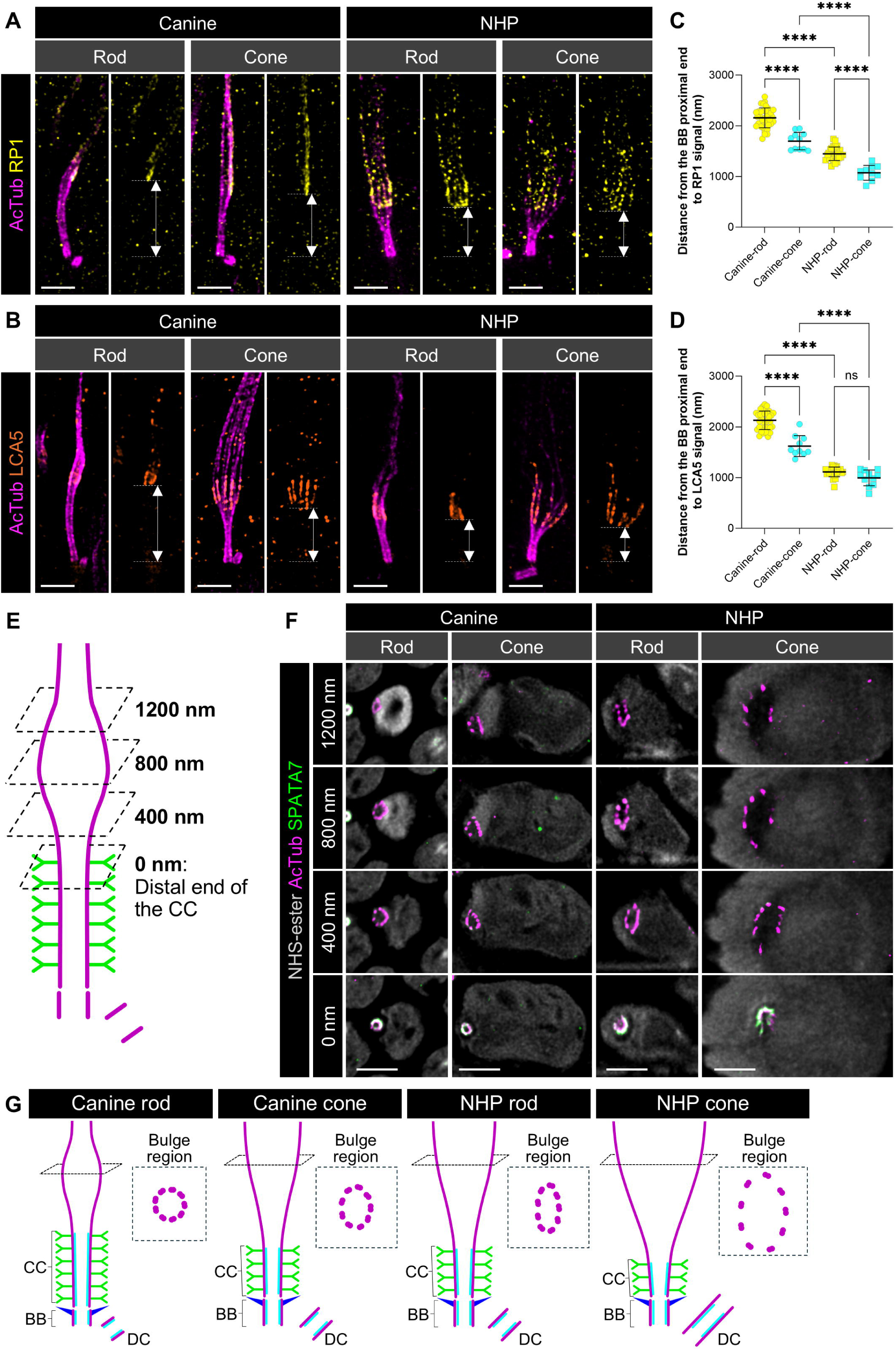
Organization of the distal region of the PSC axoneme in canine and NHP photoreceptors. **(A, B)** Representative confocal images showing the molecular architecture of the distal region of PSC axoneme in rod and cone photoreceptors from canine and NHP retinas. In addition to the AcTub-labeled axoneme (magenta), RP1 (yellow, A) and LCA5 (orange, B) were visualized by immunolabeling. White dashed lines and arrowheads indicate the distance from the proximal end of the BB to the RP1- or LCA5-positive region. All images are shown as maximum intensity projections. Scale bars, 1 μm, corrected for the expansion factor. **(C, D)** Quantification of the distance from the proximal end of the BB to RP1 (C) and LCA5 (D) signals in canine and NHP retinas (canine, circles; NHP, squares). Central lines indicate the mean, and error bars represent ± SD. Data were obtained from three canine and two NHP eyes. **** *P* < 0.0001, as assessed by Welch’s one-way ANOVA followed by Dunnett’s T3 multiple-comparison test. **(E)** Schematic diagram illustrating the imaging strategy for axial views of the distal CC/proximal OS axoneme region. The distal end of the CC was used to define the 0 nm position. **(F)** Representative axial-view images of distal ciliary axoneme in canine and NHP retinas at 0, +400, +800, and +1200 nm from the distal end of the CC. The ciliary axoneme was visualized by AcTub immunolabeling (magenta), and the connecting cilium/Y-link-associated region was visualized by SPATA7 immunolabeling (green). NHS-ester counterstaining (gray) was used as a pan-protein morphological reference to visualize surrounding photoreceptor structures. All images are shown as maximum intensity projections. Scale bars, 1 μm, corrected for the expansion factor. All microscopy images in this figure are U-ExM images. **(G)** Schematic summary diagrams illustrating structural differences between rod and cone PSCs across canine and NHP retinas. Dashed squares indicate axial views of ciliary axoneme organization in the bulge region.

Taken together, the fundamental rod–cone differences in the microtubule-based structural core were broadly conserved between canine and NHP photoreceptors: cones possessed shorter CCs, longer DCs, and a more pronounced bulge morphology than rods. At the same time, clear species-specific differences were also evident, particularly in CC length, the extent of distal CC widening, and the qualitative morphology of the bulge region (**Fig. 2G**).

### Human PSC architecture aligns with primate features while retaining rod-cone differences

Having established both shared and species-specific features of PSC architecture between canine and NHP photoreceptors, we next extended the comparison to human photoreceptors. Unlike the canine and NHP samples, which were obtained from young adult animals processed under standardized conditions, the human donor retinas were derived from elderly donors (>80 years old) and showed substantial variation in postmortem interval and fixation conditions (**Table S1**). For this reason, the human dataset was analyzed separately from the controlled canine–NHP comparison and interpreted descriptively, rather than as part of a strictly matched cross-species comparison. U-ExM analysis revealed inter-replicate variability in PSC morphology and quantitative measurements between the two human biological replicates (**Fig. 3A, B, D**). The sources of this variability remain unclear and may reflect differences in sample conditions and/or donor-specific factors. We therefore analyzed the human data both by combining measurements from the two donor eyes and by examining each biological replicate separately.

**Figure 3.**
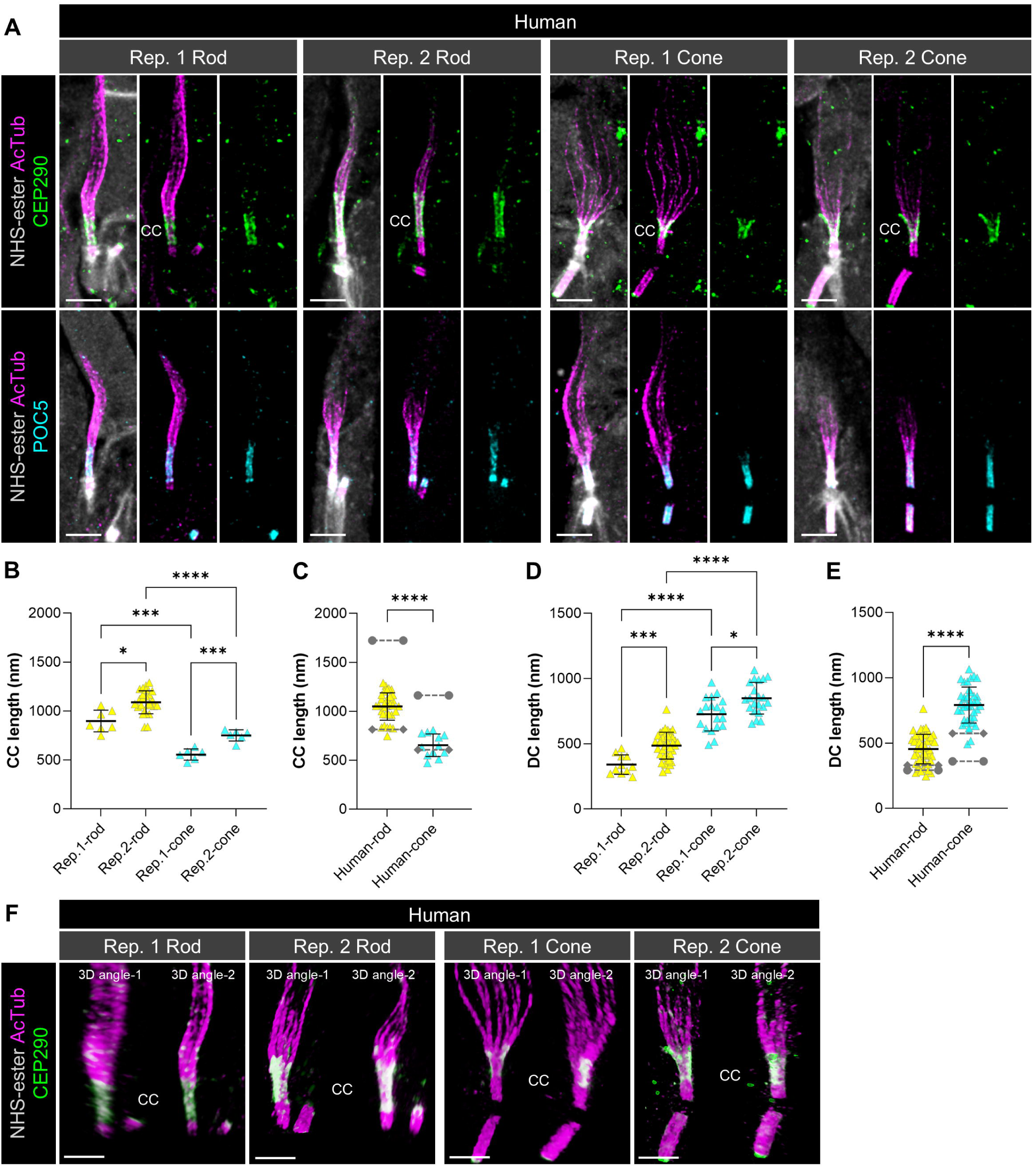
Molecular architecture of human PSCs in the context of canine and NHP PSC features. **(A)** Representative confocal images showing the ciliary axoneme visualized by AcTub immunolabeling (magenta) together with CEP290 (upper, green) and POC5 (lower, cyan) localization in human rod and cone photoreceptors from two independent biological replicates. All images are shown as maximum intensity projections. Scale bars, 1 μm, corrected for the expansion factor. **(B-E)** Quantification of CC length (B, C) and DC axoneme length (D, E). Graphs B and D show values for individual biological replicates, whereas C and E show the combined results. Gray dashed lines indicate the respective mean values for canine rods, canine cones, NHP rods and NHP cones, based on the corresponding measurements shown in Fig. 1E, J (canine, circles; NHP, squares). Central lines indicate the mean, and error bars represent ± SD. Data were obtained from two human eyes. \**P* < 0.05, *** *P* < 0.001, ** ** *P* < 0.0001, as assessed by Welch’s one-way ANOVA followed by Dunnett’s T3 multiple-comparison test (B, D) and Welch’s *t*-tests (C, E). **(F)** Three-dimensional rendered images showing differences in bulge region width from the front (angle 1) and side (angle 2) views. All microscopy images in this figure are U-ExM images.

Despite the inter-replicate variability, several fundamental rod- and cone-specific features were preserved. For CC length, significant differences were observed between the two human biological replicates, but in both samples cone CCs remained shorter than rod CCs, consistent with the findings in canine and NHP retinas (**Fig. 3B**). When the combined human data were compared descriptively with the canine and NHP datasets, human rod and cone CC lengths were shorter than the corresponding canine values, showing a pattern consistent with the shorter CC lengths observed in NHP compared with canine photoreceptors for both rods and cones (**Fig. 3C**). A similar pattern was observed for DC length. Although DC length also differed significantly between the two human biological replicates, cones consistently exhibited longer DCs than rods, again in agreement with rod-cone difference shared by canine and NHP photoreceptors (**Fig. 3D**). Notably, whereas NHP cones already displayed exceptionally long DCs, human cones exhibited even greater DC elongation than NHP cones in the combined analysis (**Fig. 3E**).

Given the striking elongation of the DCs in NHP and human cones, we next asked whether this feature differed between cone photoreceptor subtypes. Using glutamylation labeling together with cone subtype-specific opsin labeling, we measured CC and DC length in S-cones and L/M-cones across the three species (**Fig. S5A**). CC length did not significantly differ between S-cones and L/M-cones in canine, NHP or human retinas (**Fig. S5B**). In contrast, DC length was consistently greater in L/M-cones than in S-cones in all three species (**Fig. S5C**).

The distal ciliary axoneme in human photoreceptors also resembled that of NHPs more closely than that of canines. Based on qualitative morphological observations, the proximal OS axoneme/bulge region appeared more pronounced in both human rods and cones than in canine photoreceptors, and this feature was particularly evident in cones, which showed a wider axonemal bulge morphology similar to that observed in NHP cones (**Fig. 3F**). Together, these findings indicate that the molecular architecture of the microtubule-based structural core differs between canine and primate photoreceptors, whereas human photoreceptors show several structural similarities to NHP photoreceptors. At the same time, the fundamental rod–cone differences in PSC architecture were maintained across all species, with cones consistently exhibiting shorter CCs, longer DCs, and more pronounced bulge morphology than rods.

### Periciliary structures exhibit rod-cone variation and species-dependent organization in canine and NHP retinas

Previous work in primate photoreceptors has shown that the periciliary structures contain distinct Usher syndrome (USH)-associated proteins, with USH2 proteins localized to the periciliary membrane region and representative USH1 proteins associated with the IS/OS junction and calyceal process-related membranes (Sahly et al., 2012) (**Fig. 4A**). By contrast, calyceal processes are absent or vestigial in rodents, highlighting marked interspecies differences in this compartment (Sahly et al., 2012; Sharkova et al., 2023). Because the precise structural boundaries and nanoscale molecular organization of these periciliary structures remain incompletely defined, we next compared their organization between canine and NHP photoreceptors following the framework outlined above. To visualize representative components of USH1 and USH2 protein complexes, we performed immunolabeling for protocadherin-15 (PCDH15) and Whirlin, respectively, in combination with NHS-ester counterstaining, which provided a morphological reference for identifying the approximate locations of periciliary subcompartments (**Fig. 4B**).

**Figure 4.**
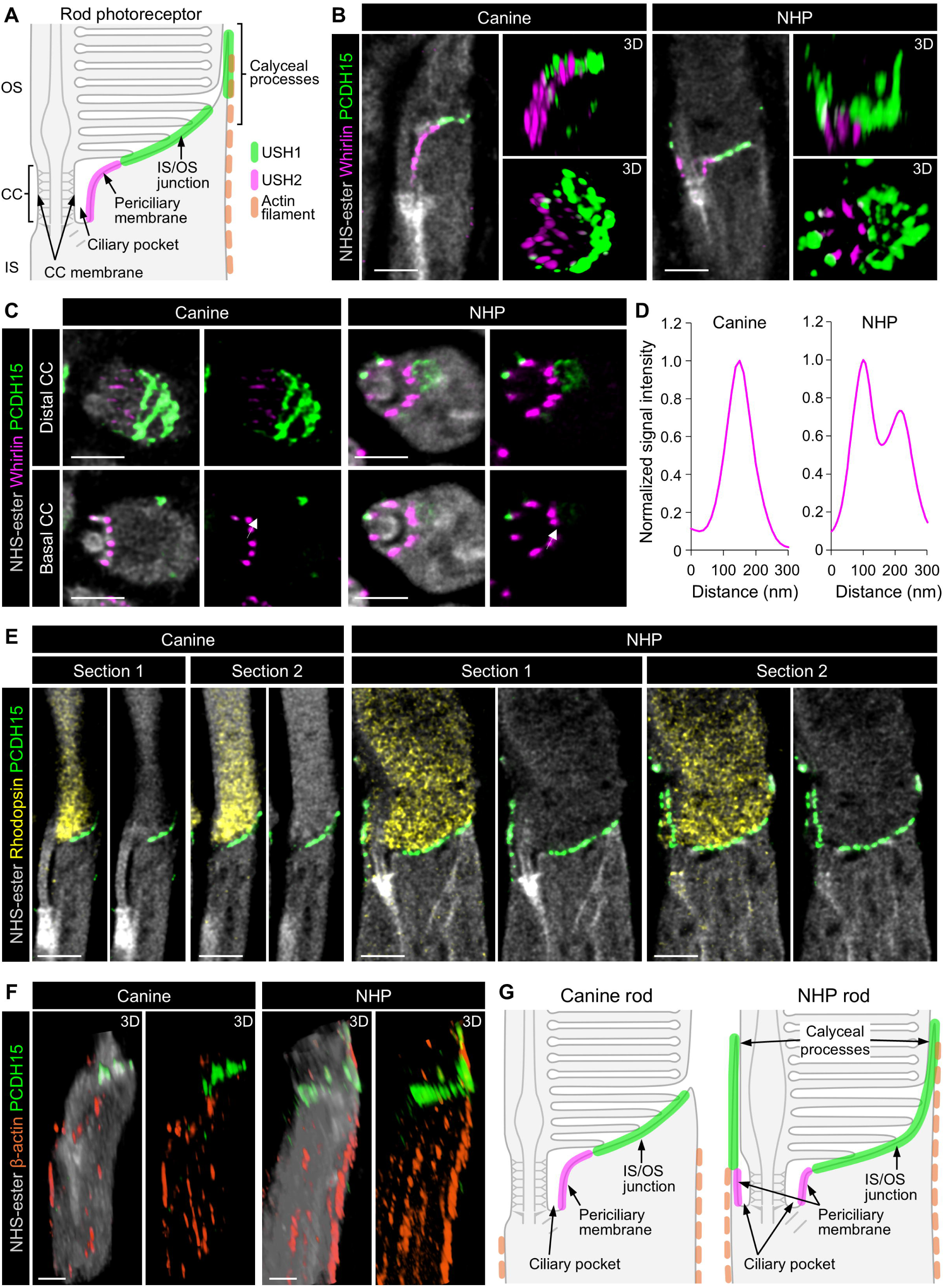
Periciliary structure organization in canine and NHP rod photoreceptors. **(A)** Schematic diagram of periciliary structures in rod PSCs based on previous publications (Sahly et al., 2012). The USH1- and USH2-associated domains shown here summarize representative localization patterns reported for selected Usher protein groups at the IS/OS junction, calyceal process, and periciliary membrane regions, and are not intended to represent all reported subcellular localizations of individual USH proteins. **(B)** Representative images showing molecular localization in periciliary structures of rod PSCs from canine and NHP retinas. Representative Usher proteins were visualized by immunolabeling for PCDH15 (USH1F, green) and Whirlin (USH2D, magenta), together with NHS-ester pan-staining (gray). For both species, left panels show single optical sections, whereas right panels show three-dimensional rendered images in lateral (upper) and axial (lower) views. Scale bars, 1 μm, corrected for the expansion factor. **(C)** Axial views of periciliary structures in rod photoreceptors corresponding to the basal (lower) and distal (upper) regions of the CC. All images are shown as single optical sections. Scale bars, 1 μm, corrected for the expansion factor. **(D)** Normalized line intensity profiles of Whirlin distribution along the periciliary membrane, corresponding to the white arrows indicated in (C). X-axis values are corrected for the expansion factor. **(E)** Single optical sections showing the distribution pattern of PCDH15 in periciliary structures of canine and NHP rods. The OS region is visualized by rhodopsin immunolabeling (yellow). For both canine and NHP images, section 1 corresponds to the region containing the CC, whereas section 2 shows the mid-peripheral region. Scale bars, 1 μm, corrected for the expansion factor. **(F)** Three-dimensional rendered images showing the distribution pattern of actin filaments associated with calyceal processes. Actin filaments are visualized by β-actin immunolabeling (orange). Scale bars, 1 μm, corrected for the expansion factor. All microscopy images in this figure are U-ExM images. **(G)** Schematic diagram illustrating the updated structural characteristics of periciliary structures in canine and NHP rod photoreceptors.

In both canine and NHP rods, Whirlin localized to the periciliary membrane domain, consistent with previous reports. However, pronounced species-dependent differences in its spatial distribution were evident. In lateral views, Whirlin signal in canine rods was restricted to the region between the CC and the IS, whereas in NHP rods it was detected on both sides of the CC. Three-dimensional reconstructions further revealed that Whirlin in canine rods covered only a restricted domain corresponding to the IS-facing side of the CC, whereas in NHP rods Whirlin-positive structures extended around the circumference of the CC (**Fig. 4B**). High-magnification axial imaging confirmed this difference in periciliary organization. NHS-ester counterstaining showed that the periciliary membrane in canine rods was confined to the IS-facing side of the CC, whereas in NHP rods it extended around the full circumference of the CC. Whirlin-positive elements were localized along these periciliary membrane domains. In NHP rods, many Whirlin-positive elements were resolved as adjacent paired pillar-like structures in at least some serial z-sections, although this paired appearance was not evident for every element in every optical section (**Fig. 4C, D; Fig. S6A**).

Similarly, the distribution of the USH1 protein PCDH15 also showed clear species-dependent differences. In canine rods, PCDH15 signal was largely restricted to the IS/OS junction region. In NHP rods, however, PCDH15 was detected not only at the IS/OS junction but also as axially oriented signals extending along the basal OS (**Fig. 4B, E**). Because calyceal processes are known to be associated with actin filaments extending from the IS, we next examined the organization of β-actin in this region. In canine rods, actin filaments within the IS appeared discontinuous and lacked clear association with PCDH15-positive structures, consistent with our previous observations (Takahashi et al., 2025). In contrast, NHP rods exhibited continuous actin filaments that were closely aligned with PCDH15 signals, forming filamentous profiles extending from the IS toward the OS base (**Fig. 4F**).

Together, these observations reveal marked species-dependent differences in the organization of rod periciliary structures. In canine rods, the periciliary membrane and ciliary pocket were restricted to the IS-facing side of the CC, whereas in NHP rods these structures extended circumferentially around the CC. Furthermore, canine rods did not show clear PCDH15/actin-associated molecular features comparable to the well-defined calyceal processes observed in NHP rods. These findings suggest that calyceal processes are absent or, if present, are very limited in canine rods. By contrast, NHP rods displayed prominent PCDH15-positive, actin-associated profiles extending along the basal OS, consistent with well-defined calyceal processes (**Fig. 4G**).

Using the same approach, we next compared the organization of periciliary structures in cone photoreceptors from canine and NHP retinas. In contrast to rods, lateral views showed Whirlin-positive elements on both sides of the CC in cones of both species. Three-dimensional reconstructions further revealed circumferential arrays of Whirlin-positive structures surrounding the CC in both canine and NHP cones, resembling the pattern observed in NHP rods (**Fig. 5A**). Notably, however, three-dimensional rendering also revealed an additional ectopic Whirlin-positive cluster in canine cones that was not associated with the periciliary membrane and was not observed in any other photoreceptor types (**Fig. 5A**). This ectopic Whirlin signal was consistently detected in all canine biological replicates examined, although its precise distribution pattern varied between cells (**Fig. S7**). High-magnification axial imaging and serial z-section analysis showed that Whirlin-positive structures were distributed circumferentially around the CC in both canine and NHP cones. Within this circumferential array, many individual Whirlin-positive puncta could be resolved into adjacent paired pillar-like elements, although this paired appearance was not evident for every element in every optical section (**Fig. 5B, C; Fig. S6B**). The ectopic Whirlin-positive cluster unique to canine cones was localized along the IS plasma membrane (**Fig. 5B**).

**Figure 5.**
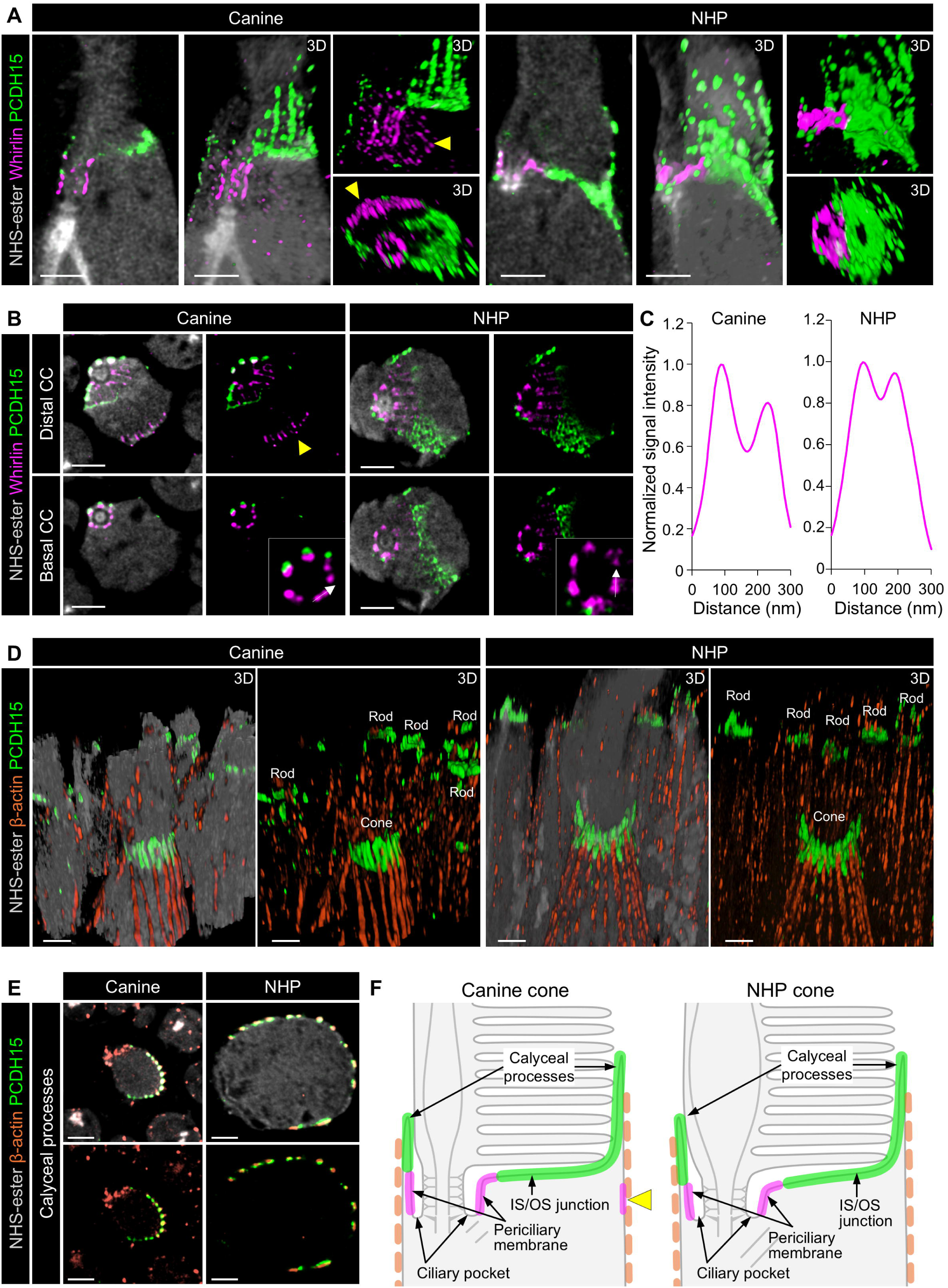
Molecular organization of periciliary structures in canine and NHP cone photoreceptors. **(A)** Representative images showing molecular localization in periciliary structures of cone PSCs from canine and NHP retinas. PCDH15 (USH1F, green) and Whirlin (USH2D, magenta) were visualized by immunolabeling, together with NHS-ester pan-staining (gray). For both species, left panels show single optical sections, whereas middle and right panels show three-dimensional rendered images in lateral and axial views. Yellow arrowheads indicate ectopic Whirlin localization in canine cones. Scale bars, 1 μm, corrected for the expansion factor. **(B)** Axial views of periciliary structures in cone photoreceptors at the basal (lower) and distal (upper) regions of the CC. Yellow arrowhead indicates ectopic Whirlin localization in canine cones. All images are shown as single optical sections. Scale bars, 1 μm, corrected for the expansion factor. **(C)** Normalized line intensity profiles of Whirlin distribution along the periciliary membrane, corresponding to the white arrows indicated in (B). X-axis values are corrected for the expansion factor. **(D)** Three-dimensional rendered images showing the distribution pattern of actin filaments associated with calyceal processes. Actin filaments are visualized by β-actin immunolabeling (orange). Scale bars, 1 μm, corrected for the expansion factor. **(E)** Axial views of calyceal processes in canine (left) and NHP (right) cone photoreceptors. All images are shown as single optical sections. Scale bars, 1 μm, corrected for the expansion factor. All microscopy images in this figure are U-ExM images. **(F)** Schematic diagram illustrating the updated structural characteristics of periciliary structures in canine and NHP cone photoreceptors.

PCDH15 exhibited a broadly similar distribution pattern in cones of both species. In canine and NHP cones, PCDH15 was detected not only at the IS/OS junction but also as axially oriented signals extending along the basal part of the OS (**Fig. 5A**). These PCDH15-positive profiles were clearly associated with actin filaments extending from the IS, supporting the presence of calyceal process in cones of both species (**Fig. 5D, E**).

Together, these findings indicate that cone photoreceptors in both canine and NHP retinas possess circumferential ciliary pocket/periciliary membrane organization around the CC, accompanied by clearly defined calyceal processes. At the same time, a species-dependent feature was evident in canine cones, which uniquely displayed an additional ectopic Whirlin-positive domain (**Fig. 5F**).

### Human periciliary structures are broadly primate-like but also show species-dependent features

We next extended the analysis of periciliary structures to human photoreceptors. In both biological replicates, rod and cone photoreceptors exhibited Whirlin-positive structures arranged circumferentially around the CC (**Fig. 6A**), consistent with the organization observed in NHP photoreceptors. In addition, both rods and cones showed PCDH15 signal not only at the IS/OS junction but also as axially oriented profiles extending along the basal OS. This axially oriented PCDH15 signal was weaker in human replicate 1 than in human replicate 2 and was more clearly appreciated in 3D-rendered images than in individual optical sections. Actin filaments extending toward these PCDH15-positive structures were also evident, further supporting a NHP-like periciliary organization in human photoreceptors (**Fig. 6B**).

**Figure 6.**
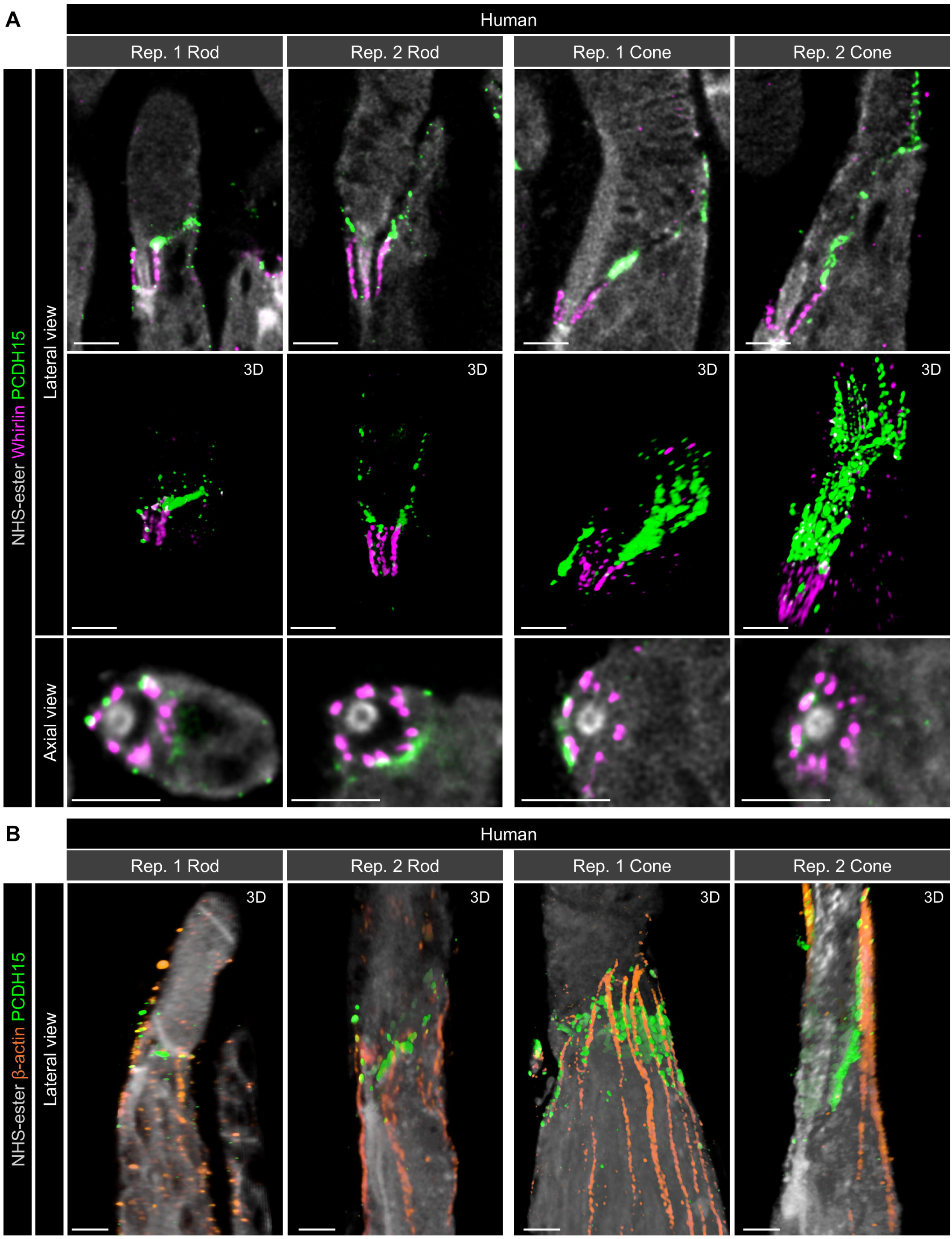
Periciliary structure architecture in human photoreceptors. **(A)** Representative images showing molecular localization in periciliary structures of rods and cones in human retina from two individual biological replicates. PCDH15 (USH1F, green) and Whirlin (USH2D, magenta) were visualized by immunolabeling, together with NHS-ester pan-staining (gray). Upper panels show single optical sections in lateral views, middle panels show three-dimensional rendered images in lateral views, and lower panels highlight axial view images at the regions of the CC. Scale bars, 1 μm, corrected for the expansion factor. **(B)** Three-dimensional rendered images showing the distribution pattern of actin filaments associated with calyceal processes. Actin filaments are visualized by β-actin immunolabeling (orange). Scale bars, 1 μm, corrected for the expansion factor. All microscopy images in this figure are U-ExM images.

Notably, the calyceal process-like structures delineated by PCDH15 and β-actin were more robust in cones than in rods (**Fig. 6B**). This rod-cone difference was consistent with the pattern observed in NHP photoreceptors and with previous reports describing more robust calyceal process organization in primate cones than in rods (Sahly et al., 2012; Verschueren et al., 2022). Together, these findings indicate that human periciliary structures broadly resemble those of NHP photoreceptors while retaining clear rod–cone differences in their structural organization.

Beyond these broadly primate-like periciliary features, we next asked whether human photoreceptors also display additional PSC-associated structures not observed in canine or NHP retinas. A recent electron microscopy-based ultrastructural study described a previously unrecognized structure unique to human rods, termed the accessory inner segment (aIS), in which the IS extends alongside the OS and is reinforced by a prominent microtubule-based scaffold. This structure was further reported to contain mitochondria extending into the OS region and to be absent from human cones (Lewis et al., 2025a). Because this finding was first reported in a recent study and has not yet been systematically evaluated in other species or by alternative imaging approaches, we used U-ExM to assess the presence and structural characteristics of aIS-like structures in canine, NHP, and human photoreceptors.

We first examined rod photoreceptors, in which the aIS has been proposed to be a human rod-specific feature (Lewis et al., 2025a). In canine and NHP rods, we found no structural features suggestive of an aIS. Consistent with a conventional rod morphology, cytochrome c-labeled mitochondria were clearly confined to conventional IS region in both species (**Fig. 7A**). Human samples, however, showed distinct findings between biological replicates. One donor retina (replicate 1) displayed rod morphology comparable to that of canine and NHP rods, whereas the other (replicate 2) contained rod profiles with aIS-like structures extending alongside the OS. In these cells, cytochrome c-positive mitochondria were also detected within the extended structure, consistent with one of the defining features of the aIS described previously. The extent of this aIS-like structure varied markedly among rods, and clear aIS-like extensions were not observed in all rod profiles in human replicate 2. Tubulin immunolabeling further showed that aIS-like structures in human replicate 2 contained α-tubulin- and β-tubulin-positive signals, but these signals were variable and uneven, resembling the broader IS microtubule network rather than forming a discrete scaffold unique to the aIS-like region (**Fig. 7A; Fig. S8A**). Consistent with the interpretation that this structure represented an IS-derived extension running alongside the OS, PCDH15 signal marked the junction between the OS and the aIS-like structure (**Fig. 7A**).

**Figure 7.**
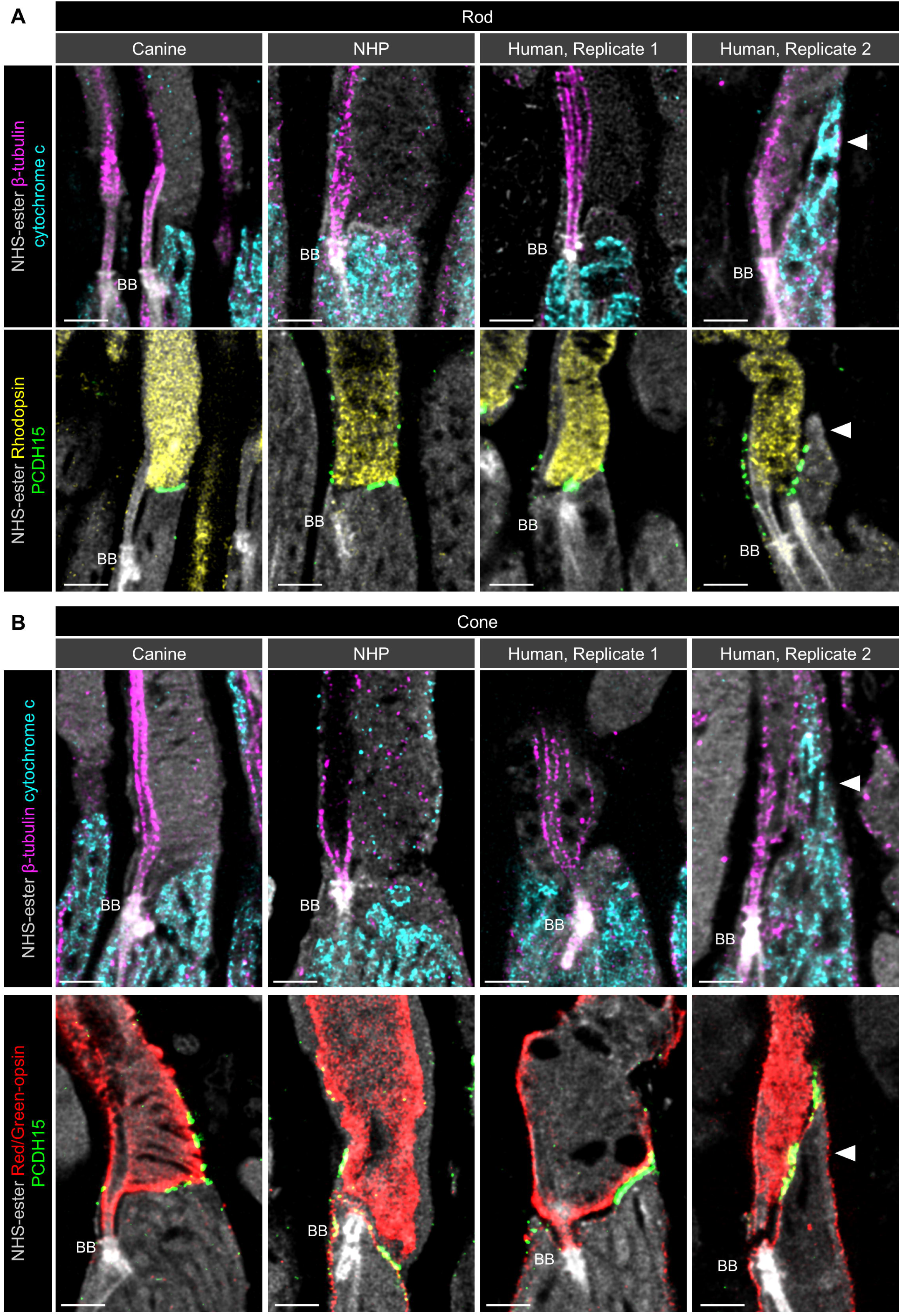
Presence or absence of the accessory inner segment across three mammalian species. **(A)** Representative single optical section images of rod photoreceptors showing the distribution of inner segment (IS) mitochondria (upper panels, cyan) and the IS/OS junction visualized by rhodopsin/PCDH15 immunolabeling (lower panels, yellow/green). Only in human replicate 2, several rod photoreceptors showed infiltration of mitochondria into the OS region and protrusion of the IS into the OS region, indicating the presence of an accessory inner segment (aIS; white arrowheads). In rod photoreceptors from canine, NHP, and human replicate 1 retinas, these structural features were not observed in any cells. **(B)** Representative images of cone photoreceptors showing the distribution of IS mitochondria (upper panels, cyan) and the IS/OS junction visualized by L/M-opsin/PCDH15 immunolabeling (lower panels, red/green). Similar to the observations in rod photoreceptors, mitochondrial infiltration into the OS region was observed in several cone photoreceptors only in human replicate 2, indicating that the aIS is also present in cone photoreceptors. All images are shown as single optical sections. Scale bars, 1 µm, corrected for the expansion factor. All microscopy images in this figure are U-ExM images.

We next performed the same analysis in cone photoreceptors. As in rods, no comparable structure was observed in canine or NHP cones. In the human samples, replicate 1 again lacked any prominent morphological variation, whereas replicate 2 contained cones with IS-derived structures extending alongside the OS (**Fig. 7B**). As in rods, these structures were supported by morphological features visualized by NHS-ester staining, the presence of cytochrome c-positive mitochondria within the OS-associated extension, tubulin-positive signals resembling the broader IS microtubule network, and an elongated PCDH15-positive IS/OS junction (**Fig. 7B; Fig. S8B**). Thus, unlike the previous report describing the aIS as a rod-specific feature of human photoreceptors (Lewis et al., 2025a), our observations suggest that aIS-like structures may also occur in human cones under at least some conditions.

Together, these findings support the possibility that aIS-like structures are a distinctive feature of human photoreceptors not observed in canine or NHP retinas. At the same time, our data indicate that their structural and molecular characteristics, their occurrence in rod versus cone photoreceptors, and their consistency across human donor samples remain unresolved. These observations therefore extend the recent ultrastructural description of the human aIS while also raising questions about its generality and rod-cone specificity.

### Species-specific and rod-cone differences extend to ciliary rootlet associated with the PSC

Finally, we assessed the morphological characteristics of the ciliary rootlet in photoreceptors across the three species. In our previous study of canine retina, we found that ciliary rootlet morphology differs strikingly between rod and cone photoreceptors. Specifically, canine rods exhibited a straight rootlet extending from the BB toward the basal IS, whereas canine cones displayed a curvilinear rootlet architecture in which the terminal portions remained connected, forming a continuous arc-like configuration within the IS (Takahashi et al., 2025). In the present dataset, canine photoreceptors showed the same rod-cone difference in rootlet morphology (**Fig. 8A,** B; Fig. S9).

**Figure 8.**
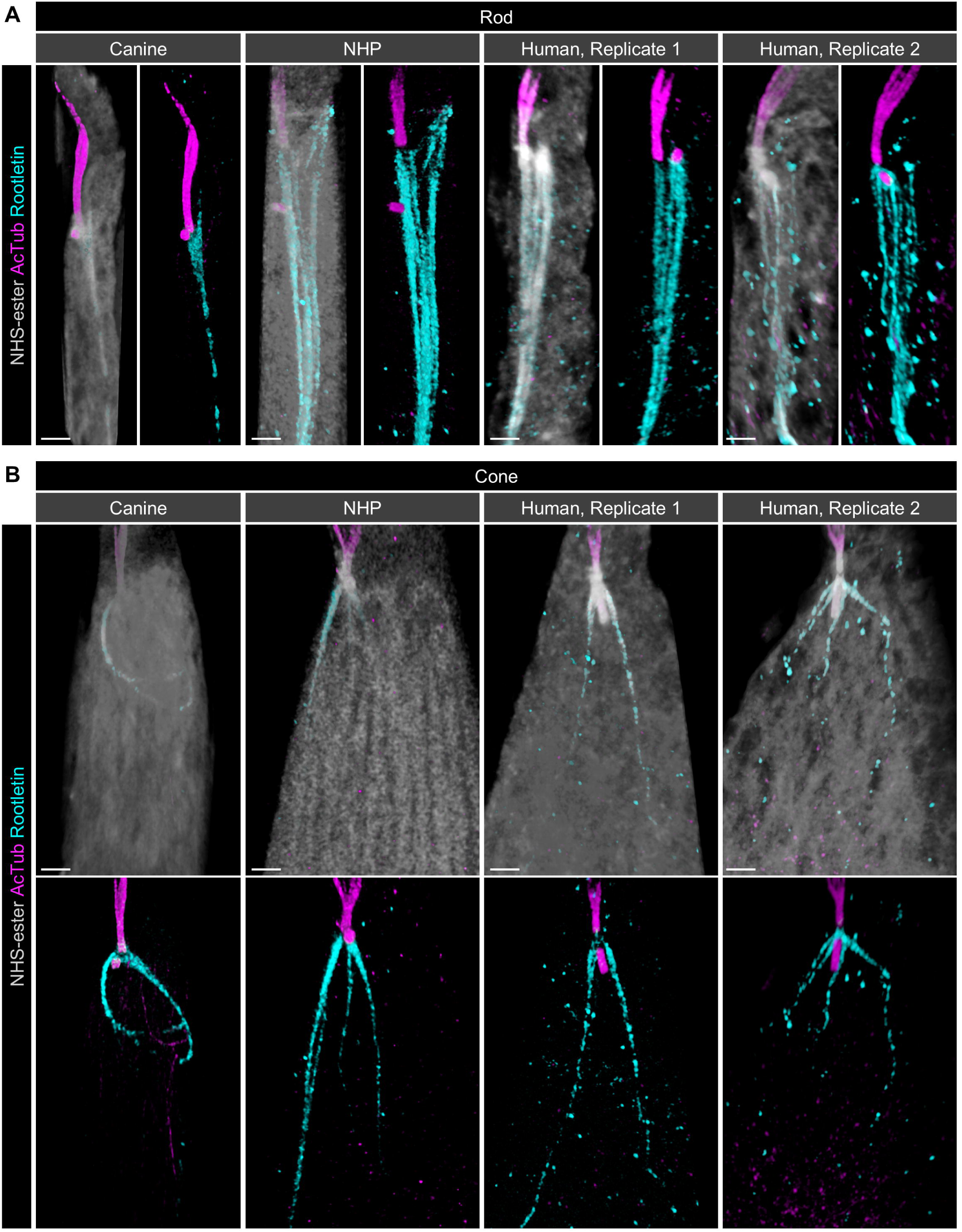
Cross-species comparison of ciliary rootlet architecture in PSCs from three mammals. **(A, B)** Representative images of rod (A) and cone (B) photoreceptors immunolabeled for AcTub (magenta) and rootletin (cyan), together with NHS-ester pan-staining (gray). NHP and human rod photoreceptors possess thicker rootlets than canine rod photoreceptors (A). Whereas the rootlet in canine cones exhibits a curvilinear morphology, NHP and human cones display rootlets that branch into two or three filaments (B). All images are shown as three-dimensional rendered images. Scale bars, 1 μm, corrected for the expansion factor. All microscopy images in this figure are U-ExM images.

By contrast, based on qualitative morphological observations of rootletin-labeled structures, ciliary rootlets in NHP and human retinas appeared to exhibit structural features distinct from those in canine retina. In rods, both NHP and human photoreceptors retained a generally straight rootlet architecture resembling that of canine rods, but these structures appeared thicker than canine rod rootlets and appeared to consist of multiple filamentous elements bundled together into a single rootlet structure (**Fig. 8A**). Cone photoreceptors showed an even more apparent species-dependent difference. Unlike canine cones, in which the rootlet terminal portions remained connected in a continuous arc-like configuration, NHP and human cones appeared to show separated rootlet termini extending as two or three thin fibers. These fibers did not appear to form a continuous arc-like terminal structure and instead appeared to extend into the intermediate region of the IS (**Fig. 8B; Fig. S9**). Thus, although the overall rod- and cone-specific configuration appeared to be preserved across species, the detailed morphology of the rootlet appeared to differ between canine and primate photoreceptors.

In contrast to the PSC axoneme and the aIS-like structure, no obvious difference in ciliary rootlet morphology was observed between the two human donor samples by qualitative morphological observation (**Fig. 8A, B**). Thus, the apparent species-specific features of human rootlets were relatively consistent across the human specimens examined. Together, these findings suggest that ciliary rootlet architecture shows clear rod–cone differences in all three species, while the specific morphological features of rod and cone rootlets appear to differ between canine and primate photoreceptors.

## Discussion

In this study, we used U-ExM to compare the nanoscale organization of the PSC and its associated structures across canine, NHP, and human retinas. Our findings reveal two broad organizational patterns. First, key rod–cone differences in the microtubule-based PSC axoneme are broadly conserved across these large mammals. Second, structures surrounding the PSC, including periciliary structures, aIS-like structures, and ciliary rootlets, exhibit substantially greater diversification among species and between rods and cones. In this context, human photoreceptors were generally more similar to those of NHPs than to those of canines, yet also displayed additional features not readily explained by the NHP pattern alone. Together, these observations refine current concepts of mammalian photoreceptor ciliary organization by showing that conserved rod- and cone-specific architectural features coexist with pronounced species-specific specialization in PSC-associated compartments.

Across all three species, cones exhibited shorter CCs, longer DCs, and more pronounced bulge morphology than rods, indicating that these rod-cone differences represent conserved architectural features of mammalian photoreceptors rather than species-restricted specialization. At the same time, primate photoreceptors differed markedly from canine photoreceptors, with shorter CCs, more pronounced widening at distal CC and bulge region, and exceptionally long cone DCs. These observations are consistent with previous mouse–human comparisons, which reported shorter CCs in human rods than in mouse rods, as well as longer DCs in human cones relative to both human rods and mouse photoreceptors (Mercey et al., 2024; Moye et al., 2025; Moye et al., 2026). Extending beyond these observations, our study demonstrates that such differences are not restricted to rodent–human divergence but are also evident across large mammalian species. Moreover, through systematic subcompartment-level analysis of the PSC in rods and cones, we demonstrate that differences in PSC organization among species and between rods and cones are more extensive and nuanced than previously appreciated. The finding that L/M-cones displayed longer DCs than S-cones across all species further suggests that PSC architecture varies not only between rods and cones, but also among cone photoreceptor subtypes. Although the functional basis of these features remains to be established, CC length and bulge region width likely reflect distinct structural requirements imposed by differences in OS organization and size between rods and cones and among species. In primate photoreceptors, the presence of larger and more elongated IS/OS profiles may favor a shorter CC as a more compact supporting segment between the IS and the enlarged OS. In parallel, the expanded bulge region in NHP and human photoreceptors may reflect the need for a more developed proximal OS axonemal region at the site where the larger OS is initiated and organized. Thus, the shorter CC and larger bulge region observed in primates may represent coordinated structural adaptations that help stabilize and organize larger photoreceptor OS compartments. The functional significance of DC elongation is more difficult to infer, because the role of the DC in mature photoreceptor cilia remains poorly understood. We therefore avoid assigning a specific mechanistic role to this feature. Nevertheless, the pronounced elongation of DCs in primate cones may represent another aspect of primate cone specialization, potentially related to the enlarged cone IS compartment and the organization of the basal ciliary apparatus.

The most conceptually important finding of this study is that the periciliary membrane is not organized uniformly across mammalian photoreceptors. In canine rods, the Whirlin-positive periciliary membrane domain was confined to the IS-facing side of the CC, whereas in primate rods and in cones of all three species it extended circumferentially around the CC. This organization implies that, in canine rods, only one side of the CC membrane is apposed to the periciliary membrane, whereas the opposite side remains relatively exposed. This configuration closely resembles the structural arrangement often inferred from electron microscopic images of mouse rods and is broadly consistent with the conventional view of rod periciliary organization (Barnes et al., 2021; May-Simera et al., 2017; Sahly et al., 2012; Spencer et al., 2019). By contrast, the circumferential Whirlin-positive domain observed in primate rods and in cones of all three species indicates that, in these photoreceptors, the CC is surrounded by the periciliary membrane and is effectively embedded within the apical IS compartment. This finding substantially revises the structural concept of the periciliary region by showing that the spatial relationship between the CC membrane and the periciliary membrane differs fundamentally across species and between rods and cones. Moreover, the presence of Whirlin-positive structures along the continuous periciliary membrane, together with differences in their arrangement among species and between rods and cones, suggests that the organization of USH2-associated domains is more complex than previously appreciated and warrants further investigation.

The circumferential Whirlin-positive organization in primate rods may also provide a structural context for the phenotypic gap between human USH2-associated retinopathy and mouse USH2 models. In humans, USH2-associated retinal disease manifests as progressive retinitis pigmentosa, with progressive visual field constriction and eventual severe visual impairment in many patients (Iannaccone et al., 2004; Pierrache et al., 2016). By contrast, mouse models affecting USH2 proteins generally show mild or very late-onset retinal degeneration compared with the human disease course (Liu et al., 2007; Yang et al., 2010; Lewis et al., 2025b). Our data suggest that this phenotypic discrepancy may partly reflect species-dependent differences in rod periciliary architecture. In primate rods, the Whirlin-positive elements surround the CC in a circumferential, cone-like manner, suggesting that the USH2-associated periciliary membrane may contribute more broadly to structural support of this region. By contrast, in rods with a more restricted periciliary domain, such as rodent and canine rods, the CC may be less dependent on USH2-associated periciliary support. This could make the structural consequences of USH2 disruption less severe or more slowly progressive in mouse models than in primate photoreceptors. This idea may also be relevant to canine inherited retinal disease (IRD) genetics. Despite the large number of naturally-occurring canine IRD models identified to date, a well-established naturally-occurring canine Usher syndrome model has not yet been described (Winkler et al., 2020). The restricted periciliary organization in canine rods raises the possibility that, as in rodents, canine rods may be relatively less vulnerable to disruption of USH2-associated periciliary support than primate rods. Although this model remains speculative and requires direct experimental testing, it provides a testable structural hypothesis linking species-specific periciliary architecture to the incomplete recapitulation of human USH2 retinopathy in non-primate animal models.

Observations of periciliary structures also refine how calyceal processes should be interpreted across species and between rods and cones. In primate rods, the combined distribution of Whirlin, PCDH15, and β-actin strongly supports the presence of well-developed periciliary membrane and calyceal processes. In contrast, canine rods showed PCDH15 localization largely restricted to the IS/OS junction, limited PCDH15–actin association, and a unilateral periciliary membrane organization, suggesting either absence or very limited development of calyceal process-like structures in this photoreceptor population. This pattern closely resembles the rod phenotype previously described in mice and suggests that limited rod calyceal specialization may extend beyond rodents (Sahly et al., 2012). However, canine cones displayed a clear calyceal process-like molecular organization, although, to our knowledge, calyceal processes in canine cones have not been definitively documented by prior electron microscopy studies. This rod–cone distinction in the canine retina raises the possibility that species currently regarded as calyceal process-negative may still harbor cone-specific or rudimentary periciliary specializations that have been overlooked. Although mouse photoreceptors are generally considered to lack typical calyceal processes, the possibility of cone-restricted calyceal process-like specializations has been raised in a limited prior study, warranting targeted re-evaluation with rod- and cone-resolved structural and molecular analyses (Yang et al., 2010).

Human photoreceptors were broadly NHP-like in their periciliary organization, although the human samples also highlighted the difficulty of generalizing from a limited number of donor eyes with variable sample conditions. Both human replicates showed circumferential Whirlin-positive arrays and robust PCDH15/actin-defined calyceal process-like structures in rods and cones, supporting the conclusion that human periciliary structures are fundamentally primate-like. By contrast, aIS-like structures were highly variable. They were absent from both eyes in the non-macular pericentral region of donor 1, whereas donor 2 contained variable rod profiles with aIS-like extensions. In donor 2, these aIS-like extensions were not observed in all rod profiles, and their morphology varied from relatively long OS-apposed extensions to more limited profiles. Because the prevalence of aIS-positive rods and the extent of cell-to-cell morphological variability were not defined in the previous study (Lewis et al., 2025a), further analysis will be required to determine how consistently this architecture is present in human rods. We also observed aIS-like extensions in cones, which differs from a recent study that emphasized the aIS as a human rod-specific architecture. Thus, our data support the idea that aIS-like structures may represent a distinctive feature of human photoreceptors, but they also indicate that their prevalence, photoreceptor population specificity, consistency across human donors, and molecular definition remain unresolved and require validation in larger human cohorts.

An important consideration is retinal topography. In this study, we did not analyze the macular or foveal region in primate and human retinas, nor did we specifically examine the canine area centralis/fovea-like region. This is relevant because foveal and perifoveal cones are known to differ from more peripheral cones in density and morphology, including elongation of cone OS (Tschulakow et al., 2018; Westheimer, 2008). Similarly, the canine retina contains a specialized area centralis with a primate fovea-like cone bouquet, characterized by high cone density and elongated IS and OS despite the absence of a true foveal pit (Beltran et al., 2014). Therefore, our conclusions should be interpreted as applying primarily to the sampled superior central, non-foveal retinal regions. Future rod- and cone-resolved U-ExM analysis of the primate fovea and canine area centralis will be needed to determine whether cones in these specialized retinal regions exhibit additional PSC, periciliary membrane, calyceal process, or aIS-like specializations.

Several limitations should be considered when interpreting these findings. Most importantly, the human donor tissues differed substantially from the canine and NHP samples in age, postmortem interval, and fixation conditions, and therefore do not constitute a strictly matched cross-species comparison. In addition, U-ExM provides nanoscale molecular information but does not directly replace electron microscopy for defining ultrastructural boundaries. Accordingly, some of our morphological interpretations, particularly for calyceal process-like and aIS-like structures, remain inference-based at the molecular level. Another technical consideration is that dimensional measurements obtained from U-ExM images depend on expansion factor correction. Therefore, EF-corrected values should be interpreted primarily as relative measurements among samples processed and imaged under standardized conditions, rather than as direct substitutes for measurements obtained from non-expanded specimens. Despite these limitations, the consistency of multiple markers across species and between rods and cones supports the robustness of the major organizational patterns identified here. Overall, our study provides a comparative nanoscale framework for mammalian photoreceptor ciliary organization and shows that the strongest diversification occurs not in the core PSC axoneme itself, but in the associated membrane and support structures that surround it. These findings substantially advance current understanding of photoreceptor ciliary architecture by revealing how conserved structural features coexist with species-specific specializations and rod–cone differences.

## Materials and Methods

### Animals

Cryopreserved archival retinal tissues from adult dogs (*n* = 3; 24 weeks old) and cynomolgus macaques (*Macaca fascicularis*; *n* = 2; 4 years old) were used in this study. These samples were not collected specifically for the present work, but were residual archival materials from previous studies and were reused here for comparative ultrastructural analysis; details of animal husbandry and tissue collection have been described previously (Jacobson et al., 2007; Takahashi et al., 2026). All procedures conformed to the ARVO Statement for the Use of Animals in Ophthalmic and Vision Research and were approved by the Institutional Animal Care and Use Committee of the University of Pennsylvania. Details of the individual retinal samples used in this study are provided in **Table S1**.

### Human donor eye tissue

Enucleated human donor eyes were obtained from the San Diego Eye Bank. The use of postmortem human eyes was exempted of ethical approval by Institutional Review Boards at the University of Pennsylvania, yet all procedures were conducted in accordance with the university safety guidelines and regulations. Retinas used in this study were derived from an 83-year-old male donor (replicate 1) and an 84-year-old female donor (replicate 2). Only donors without ocular pathology were included. All tissues were de-identified before delivery to the laboratory. Details of the donor samples are summarized in **Table S1**.

### Retinal tissue processing

Procedures for canine and NHP ocular tissues were performed as described previously (Jacobson et al., 2007; Takahashi et al., 2024). The fixation regimen was standardized across all canine and NHP specimens, consisting of 4% formaldehyde for 3 h followed by 2% formaldehyde for 24 h. After cryoprotection by sequential incubation in 15% and 30% sucrose solutions in 1× phosphate-buffered saline (PBS), all formaldehyde-fixed retinal tissues were embedded in optimal cutting temperature (OCT) compound, snap-frozen in liquid nitrogen and stored at −80°C. Frozen OCT blocks were cryosectioned at 20 μm, and the sections were stored at −20°C until further use.

Human donor retinal tissues were processed under conditions that differed from those used for canine and NHP tissues. In particular, the two human donor samples were processed under distinct postmortem and fixation conditions: replicate 1 was fixed approximately 3 h after death and remained in 4% formaldehyde for 7 days, whereas replicate 2 was fixed approximately 12 h after death and remained in 4% formaldehyde for 24 h. After fixation, human tissues were cryoprotected, embedded in OCT compound, frozen and cryosectioned as described above. Details of the procedures for individual samples are summarized in **Table S1**.

### Lectin Staining on Non-Expanded Retinal Sections

Frozen OCT blocks were cryosectioned at a thickness of 10 µm and stored at −20°C until further processing. Sections were thawed at room temperature for 30 minutes, washed with PBS to remove residual OCT compound, and incubated with fluorescein-conjugated wheat germ agglutinin (WGA) and peanut agglutinin (PNA) in PBS for 30 minutes at room temperature without prior blocking. After incubation, the sections were washed with PBS and mounted for imaging. The lectins used are listed in **Table S3**.

### Ultrastructure Expansion Microscopy (U-ExM) on retinal cryosections

Retinal cryosections were processed using a U-ExM workflow previously established for long-term cryopreserved retinal tissues (Takahashi et al., 2026; Takahashi et al., 2025). Briefly, 20-µm-thick cryosections were thawed and washed with PBS to remove OCT compound, followed by overnight incubation in an anchoring solution containing 2% acrylamide and 1.4% formaldehyde. The sections were then incubated in monomer solution composed of 23% (w/v) sodium acrylate, 10% (w/v) acrylamide, 0.1% (w/v) N,N′-methylenebisacrylamide and 1× PBS. Gelation was performed under a coverslip using monomer solution supplemented with 0.5% ammonium persulfate and 0.5% tetramethylethylenediamine. Following gelation, samples were denatured at 95°C in buffer containing 200 mM sodium dodecyl sulfate (pH = 9) and subsequently expanded in distilled water for the first-round expansion. After first-round expansion, gels were equilibrated in 1× PBS and then incubated in blocking buffer for 30 minutes. The gels were incubated with primary antibodies diluted in blocking buffer overnight, followed by incubation with fluorophore-conjugated secondary antibodies diluted in blocking buffer for 2 hours at room temperature. When indicated, NHS-ester staining was performed after immunolabeling as a pan-protein counterstain. After the labeling procedure, gels were re-expanded in excess distilled water for the second-round expansion and kept at 4°C until imaging. Reagents used for U-ExM and fluorescence labeling are listed in **Tables S2 and S3**.

### Confocal imaging and image processing

Images were acquired using a STELLARIS 8 confocal microscope (Leica Microsystems, Wetzlar, Germany), as described previously (Takahashi et al., 2025). U-ExM samples were imaged using the 63× water-immersion objective (NA 1.20). For imaging, expanded gels were transferred to poly-D-lysine-coated glass-bottom dishes. The dishes were kept humidified during image acquisition to prevent gel drying and shrinkage, and imaging of each gel was completed within 30 minutes. For lateral views, z-stacks were acquired at 0.4 μm intervals with an *x,y* pixel size of 45.09 nm and a scan speed of 400 Hz. Axial-view images were acquired with an *x,y* pixel size of 22.55 nm and a scan speed of 100 or 200 Hz. Non-expanded retinal sections were imaged using the 100× oil-immersion objective (NA 1.40). Representative and quantitative images were processed in Leica Application Suite X (LAS X; Leica Microsystems) for LIGHTNING deconvolution with default parameters, as well as for generation of merged images and maximum-intensity projections. Three-dimensional (3D) rendering was performed using Imaris software (version 10.2.0; Oxford Instruments, Bristol, UK) in blend mode.

### Quantification

#### Expansion factor (EF) calculation

The EF was estimated by dividing the mantuary-measured width of the acetylated α-tubulin (AcTub)-labeled basal body (BB) axoneme in U-ExM-processed canine and NHP samples by the reported width of the non-expanded centriole in human U2OS cells (231.3 nm), assuming conservation of BB (mother centriole) width across mammalian species (Faber et al., 2023; Mercey et al., 2024; Mercey et al., 2022). Because BB axoneme width was comparable between canine and NHP retinas, a single EF of 3.64 was determined by averaging the values obtained from canine and NHP photoreceptors. This EF was applied to all quantitative measurements.

#### Protein signal length, width, and distance

Protein signal lengths, widths and distances from the proximal end of the BB were measured manually in the corresponding regions using the “Draw scalebar” function in LAS X software, as described previously (Takahashi et al., 2025). All measurements were subsequently corrected using the EF.

#### Line-profile analysis

Intensity profiles were extracted using the “Quantify” tool in LAS X. The exported intensity and distance values were imported into Microsoft Excel, and distance values were corrected by the EF. Intensities were normalized for each channel such that the maximum value within the measured range was set to 1. Line-profile plots were generated in Microsoft Excel.

### Statistical analysis

Pairwise comparisons were performed using Welch’s *t*-test, whereas comparisons involving more than two groups were analyzed using Welch’s one-way ANOVA followed by Dunnett’s T3 multiple-comparison test. Quantitative data are presented as scatter dot plots, with central lines and error bars indicating the mean and standard deviation (± SD), respectively. Statistical significance was defined as follows: ns, not significant (*P* > 0.05); * *P* < 0.05, ** *P* < 0.01, *** *P* < 0.001, and **** *P* < 0.0001. All statistical analyses were performed using Prism 10 (GraphPad, Boston, MA, USA). Source data for all graphs are provided in the **Supplementary Data**.

## Data availability

All numerical source data underlying the quantitative analyses are included in the **Supplementary Data**. Representative images are shown in the main figures and supplementary figures. Source images are available from the corresponding authors upon reasonable request.

## Author Contributions

K.T. and W.A.B. participated in the conception, experimental design, and interpretation of the data for this work. K.T. and M.O. conducted data acquisition and quantitative analysis of data. K.T. is responsible for statistical analysis and the drafting of the manuscript. K.T., M.O., R.S., and W.A.B. reviewed and edited the manuscript. All authors reviewed the manuscript, approved its submission, and take full responsibility for the manuscript.

## Acknowledgements

We express our gratitude to Drs. Qin Liu and Eric A. Pierce of Harvard Medical School for providing the RP1 antibody (Liu et al., 2002); Dr. Gordon Ruthel of Penn Vet Imaging Core Facility (Research Resource Identifier: SCR_022764) for technical support; the Division of Experimental Retinal Therapies staff for their excellent support. During the preparation of this manuscript, we used ChatGPT Plus (version 5.4) to improve language and readability. After using this tool, we reviewed and edited the content as needed and take full responsibility for the content of the publication.

