## Supplemental Figures and Tables for "Cross-species molecular mapping of the photoreceptor sensory cilium and periciliary structures identifies conserved and species-specific architectural features"

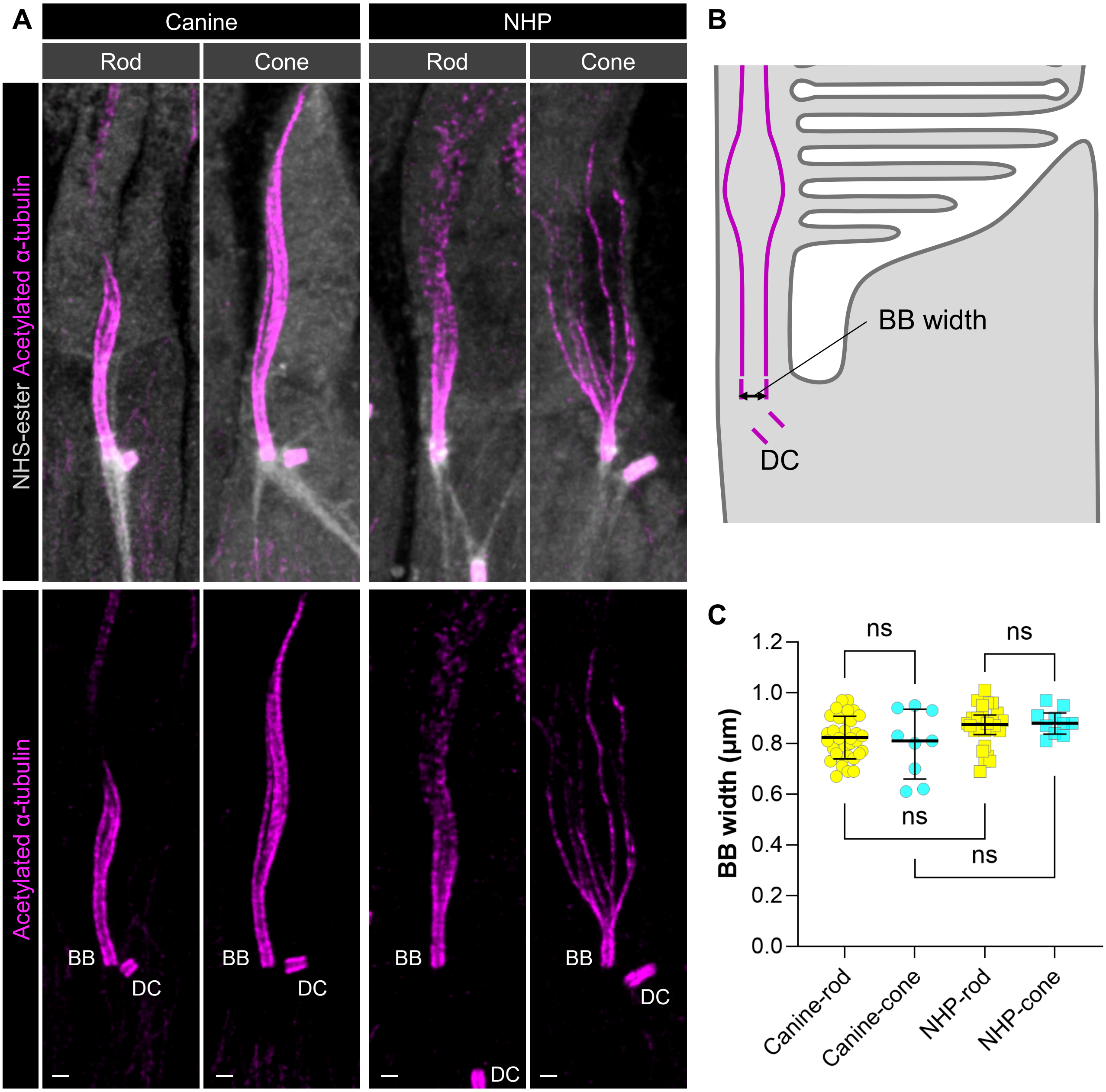


**Fig. S1. Comparison of basal body width for determination of the expansion factor.**

**(A)** Representative confocal images of photoreceptor sensory cilium (PSC) architecture in rod and cone photoreceptors from canine and non-human primate (NHP) retinas. PSC axoneme was visualized by acetylated α-tubulin (AcTub) immunolabeling (magenta). N-hydroxysuccinimide (NHS) -ester counterstaining (gray) was used as a pan-protein morphological reference to visualize surrounding photoreceptor structures. All images are shown as maximum intensity projections. Scale bars, 1 μm, shown without correction for the expansion factor. All microscopy images in this figure are U-ExM images. **(B)** Schematic diagram illustrating basal body (BB) width measurement. **(C)** Cross-species comparison of BB width between rods and cones. Central lines indicate the mean, and error bars represent ± SD. No significant differences were observed among any pairwise comparisons (Welch's one-way ANOVA followed by Dunnett's T3 multiple-comparison test; n.s., $P > 0.05$).

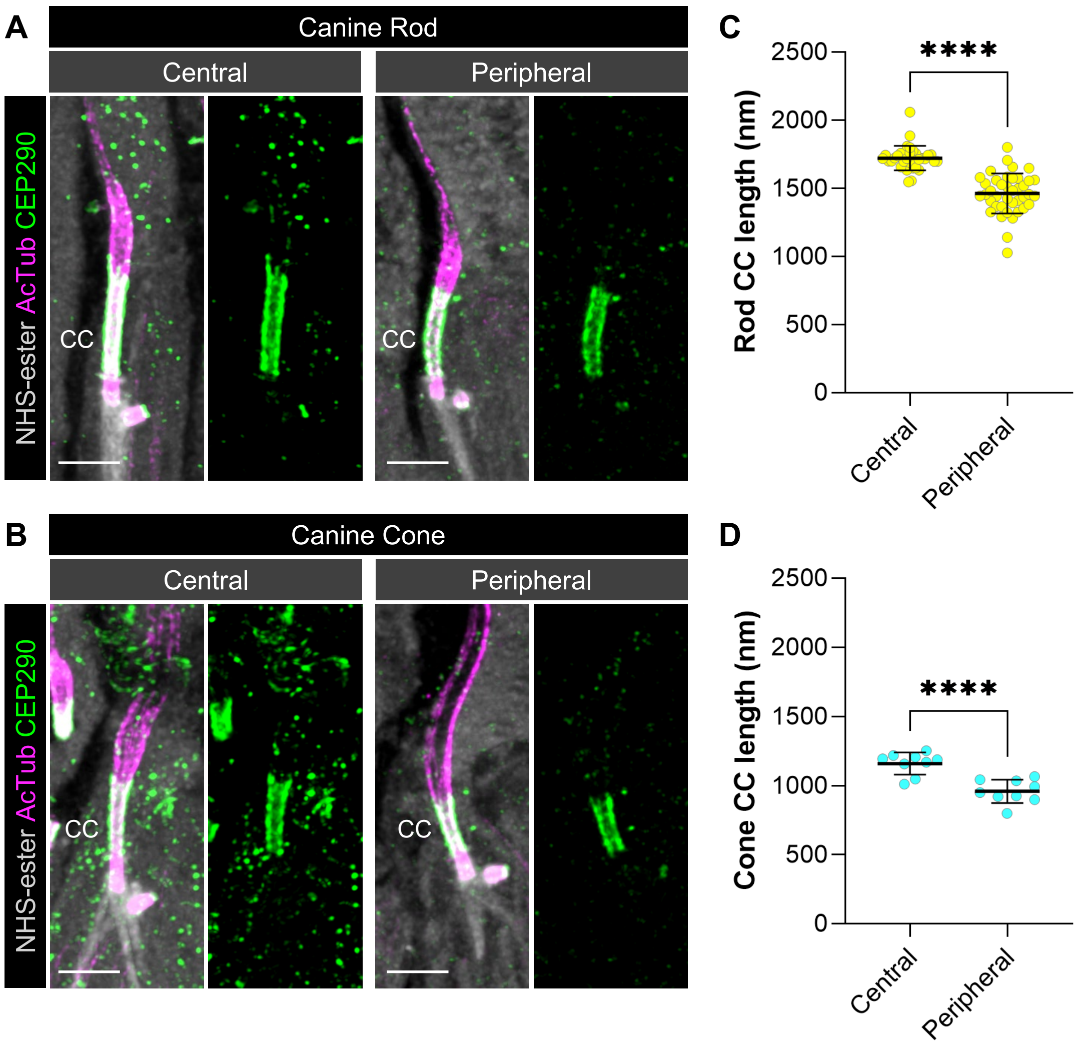


**Fig. S2. Comparison of CC length between central and peripheral regions in the canine retina.**

**(A, B)** Confocal images showing the length of the connecting cilium (CC) in the central and peripheral regions of the canine retina for rods (A) and cones (B). The ciliary axoneme and the CC were visualized by AcTub/CEP290 immunolabeling (magenta/green). NHS-ester counterstaining (gray) was used as a pan-protein morphological reference to visualize surrounding photoreceptor structures. All images are shown as maximum intensity projections. Scale bars, 1 μm, corrected for the expansion factor. All microscopy images in this figure are U-ExM images. **(C, D)** Quantification of CC length in the central and peripheral retina for rods (C) and cones (D). Central lines indicate the mean, and error bars represent ± SD. ****$P$ < 0.0001, as assessed by Welch's $t$-test.

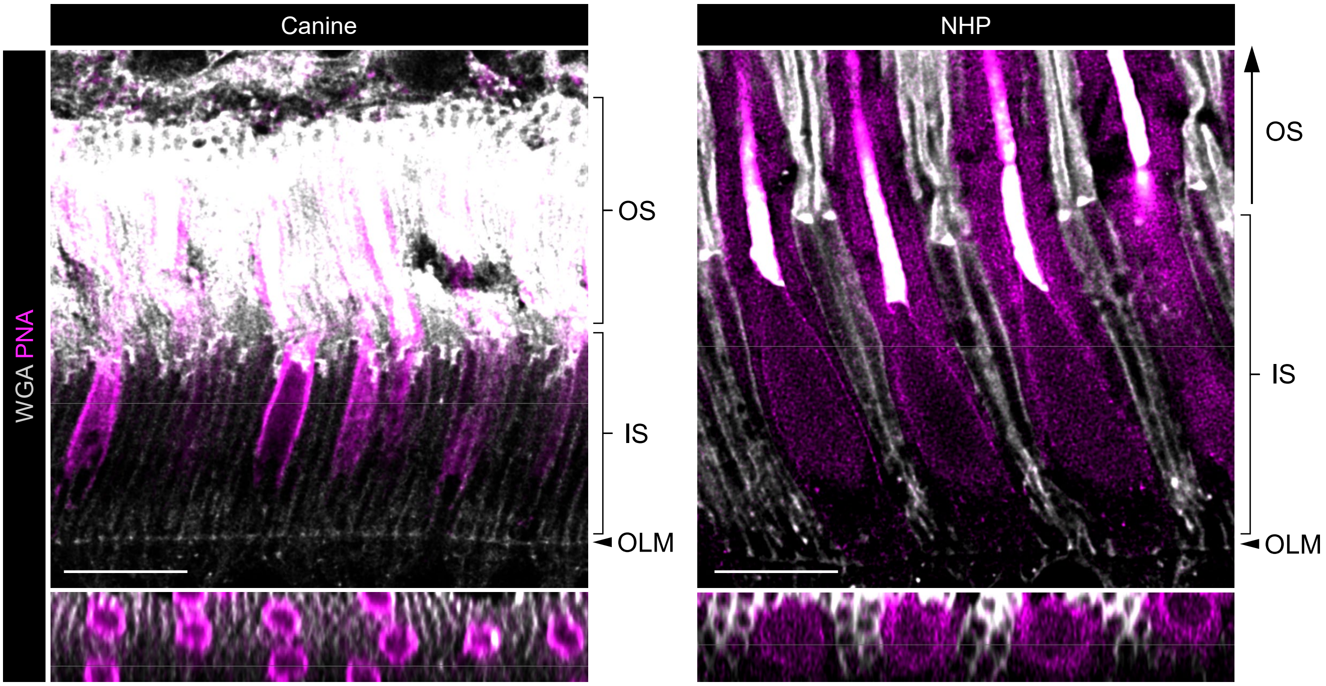


**Fig. S3. Non-expanded retinal sections showing photoreceptor IS/OS morphology in canine and NHP retinas.**

Representative single z-slice confocal images of non-expanded canine and NHP retinal sections stained with wheat germ agglutinin (WGA; gray) and peanut agglutinin (PNA; magenta). The field of view includes the outer limiting membrane (OLM), inner segment (IS), and outer segment (OS) regions, corresponding to the region shown in Fig. 1B. Lower panels show transverse views through the IS region. These images show that NHP photoreceptors have longer IS/OS profiles and greater transverse width at the IS level than canine photoreceptors. Scale bars, 10 μm.

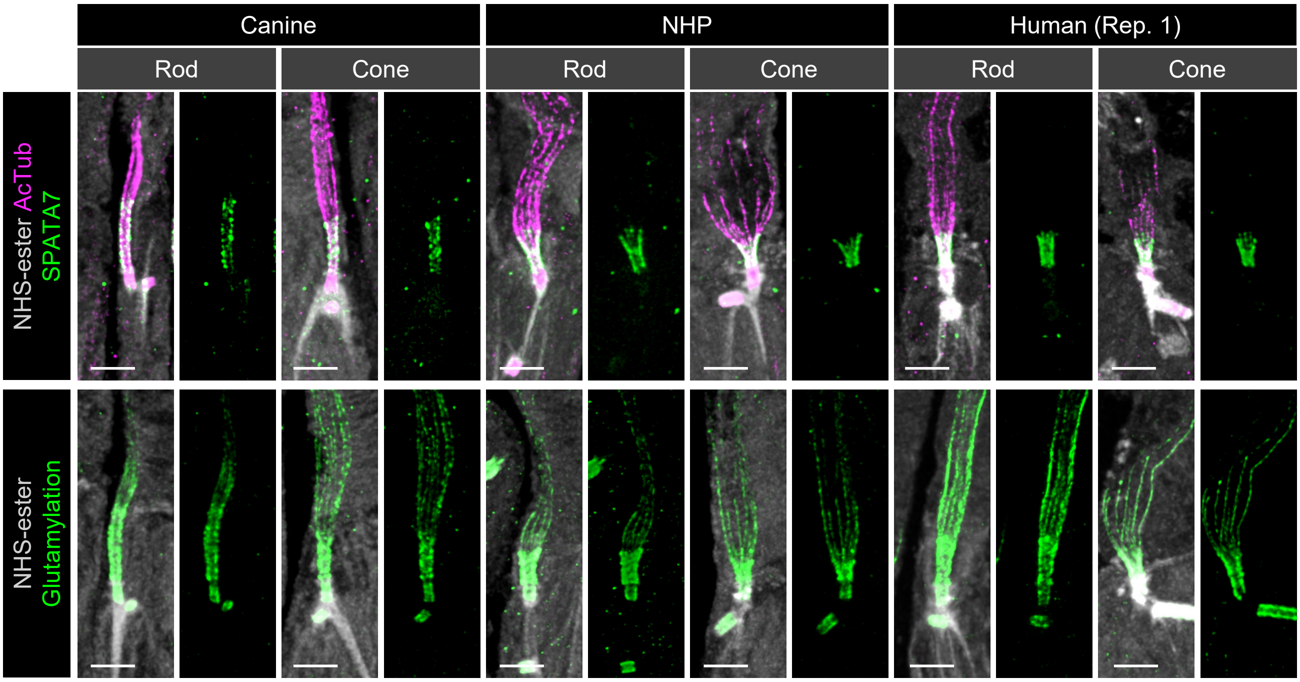


**Fig. S4. Representative molecular markers of the CC.**

Representative confocal images showing additional molecular markers of the microtubule-based structural core and CC length in rod and cone photoreceptors from canine, NHP, and human retinas. Upper panels show AcTub (magenta) and SPATA7 (green) immunolabeling, whereas lower panels show glutamylation (green) labeling of the PSC. Consistent with the results shown in Figs. 1 and 3, NHP and human photoreceptors have shorter CCs than canine photoreceptors, whereas the difference in CC length between rods and cones is maintained across all species. In addition, NHP and human cone photoreceptors possess markedly longer DCs than those in their rod counterparts and in canine photoreceptors. All images are shown as maximum intensity projections. Scale bars, 1 µm, corrected for the expansion factor. All microscopy images in this figure are U-ExM images.

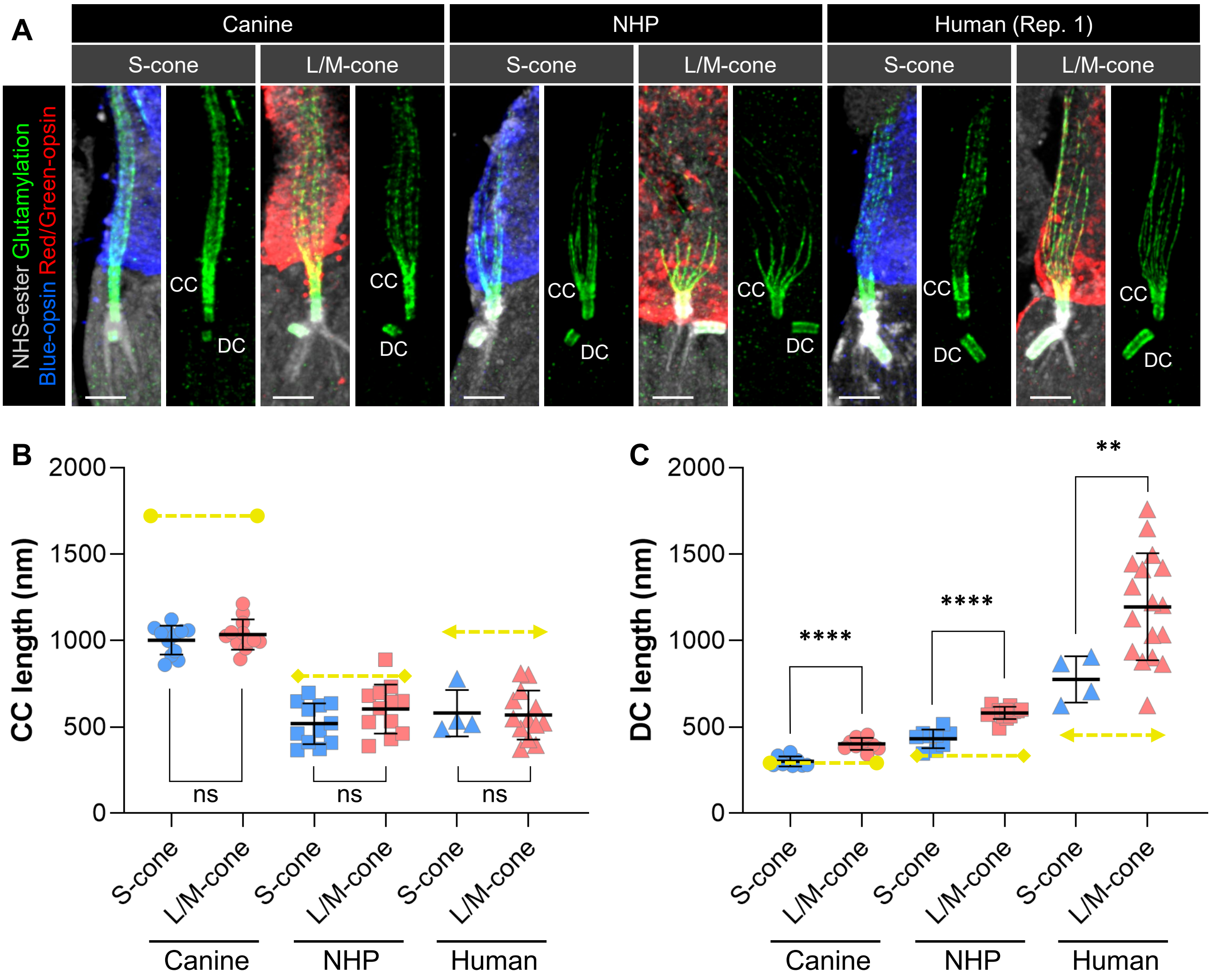


**Fig. S5. Comparison of CC and DC length between S-cones and L/M-cones.**

**(A)** Confocal images showing PSC structures in canine, NHP, and human retinas for S-cones (blue) and L/M-cones (red). The ciliary axoneme and CC were visualized by glutamylation immunolabeling (green). All images are shown as maximum intensity projections. Scale bars, 1 μm, corrected for the expansion factor. All microscopy images in this figure are U-ExM images. **(B, C)** Quantification of CC length (B) and DC length (C) in the three species (canine, circles; NHP, squares; human, triangles). Central lines indicate the mean, and error bars represent the ± SD. Yellow dashed lines indicate the mean values for rod photoreceptors of each species, derived from the corresponding measurements shown in Fig. 1E, J and Fig. 3C, E. Data were obtained from two or three individual eyes for each species. $^{**}P < 0.01$, $^{****}P < 0.0001$, as assessed by Welch's *t*-test.

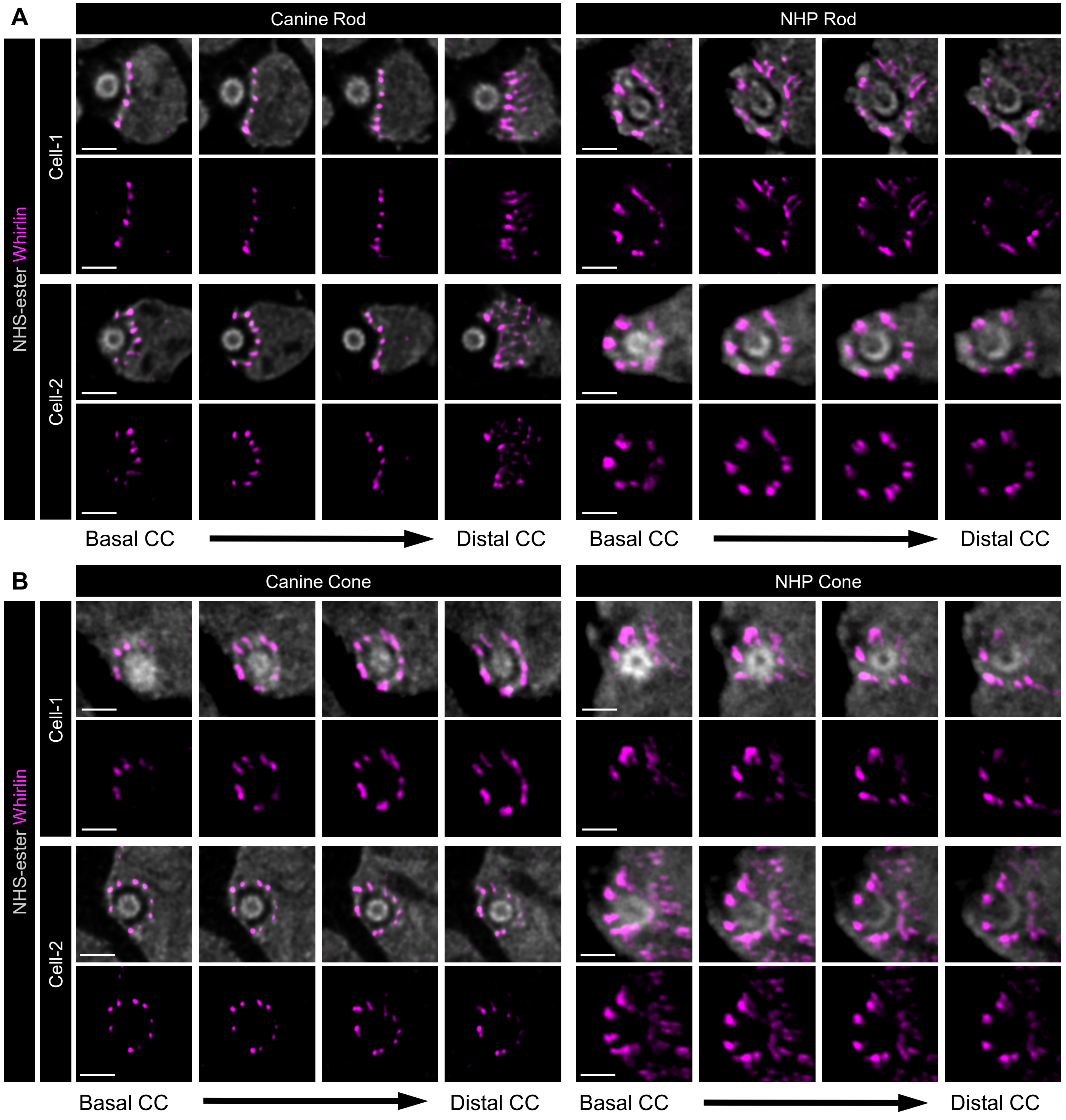


**Fig. S6. Serial z-section analysis of Whirlin-positive periciliary structures in canine and NHP photoreceptors.**

**(A, B)** Representative serial axial-view U-ExM images through the CC/periciliary region are shown for rods (A) and cones (B). Two representative photoreceptors are shown for each group: canine rods, NHP rods, canine cones, and NHP cones. In canine rods, Whirlin-positive structures were restricted to the IS-facing side of the CC, whereas in NHP rods, canine cones, and NHP cones, Whirlin-positive structures were distributed circumferentially around the CC. Many individual Whirlin-positive elements could be resolved as adjacent paired pillar-like structures in at least some z-sections, although this paired appearance was not evident for every element in every optical section. Scale bars, 500 nm, corrected for the expansion factor. All microscopy images in this figure are U-ExM images.

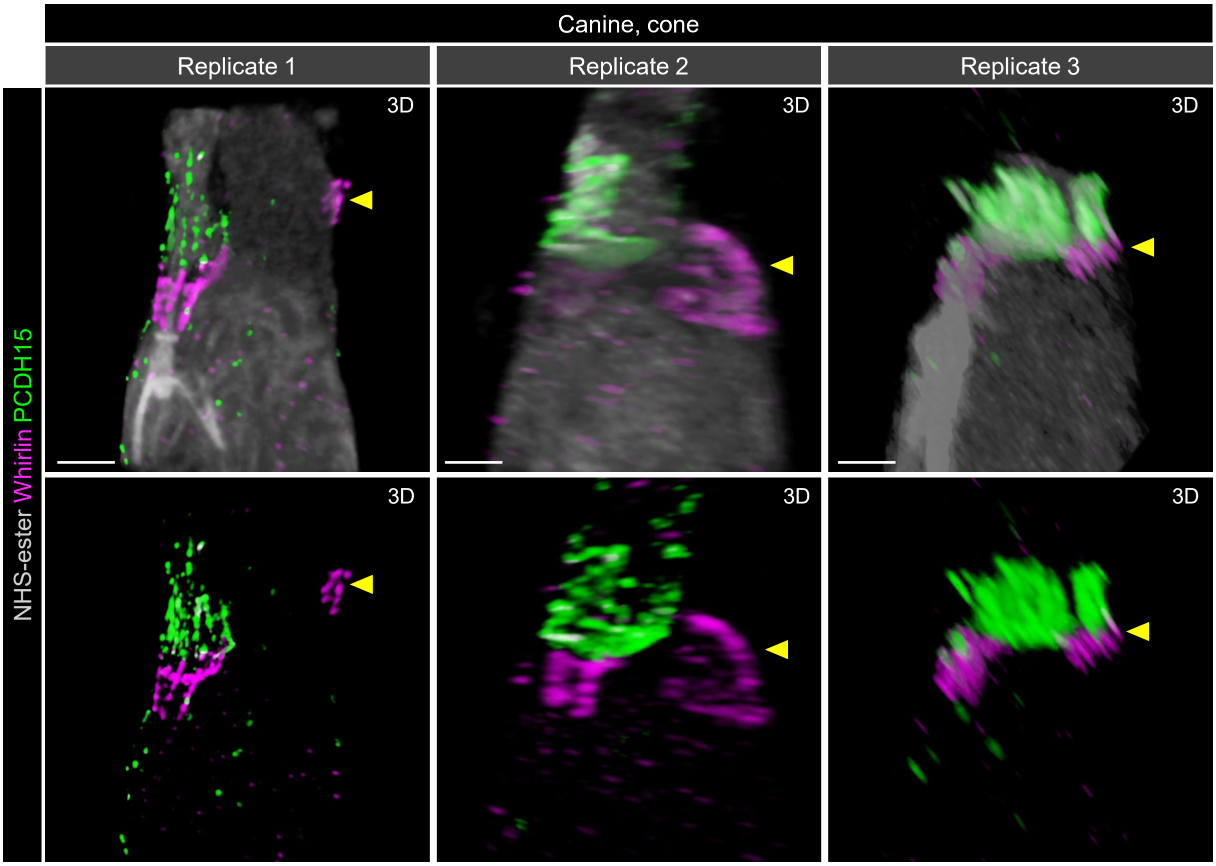


**Fig. S7. Ectopic localization of Whirlin in canine cone photoreceptors.**

Representative confocal images showing the localization of whirlin (magenta) and PCDH15 (green) in periciliary structures of canine cone photoreceptors from three independent biological replicates. Upper panels show merged images of whirlin, PCDH15, and NHS-ester counterstaining (gray), whereas lower panels show whirlin and PCDH15 labeling only. Yellow arrowheads indicate ectopic localization of whirlin. All images are shown as 3D-rendered images. Scale bars, 1 μm, corrected for the expansion factor. All microscopy images in this figure are U-ExM images.

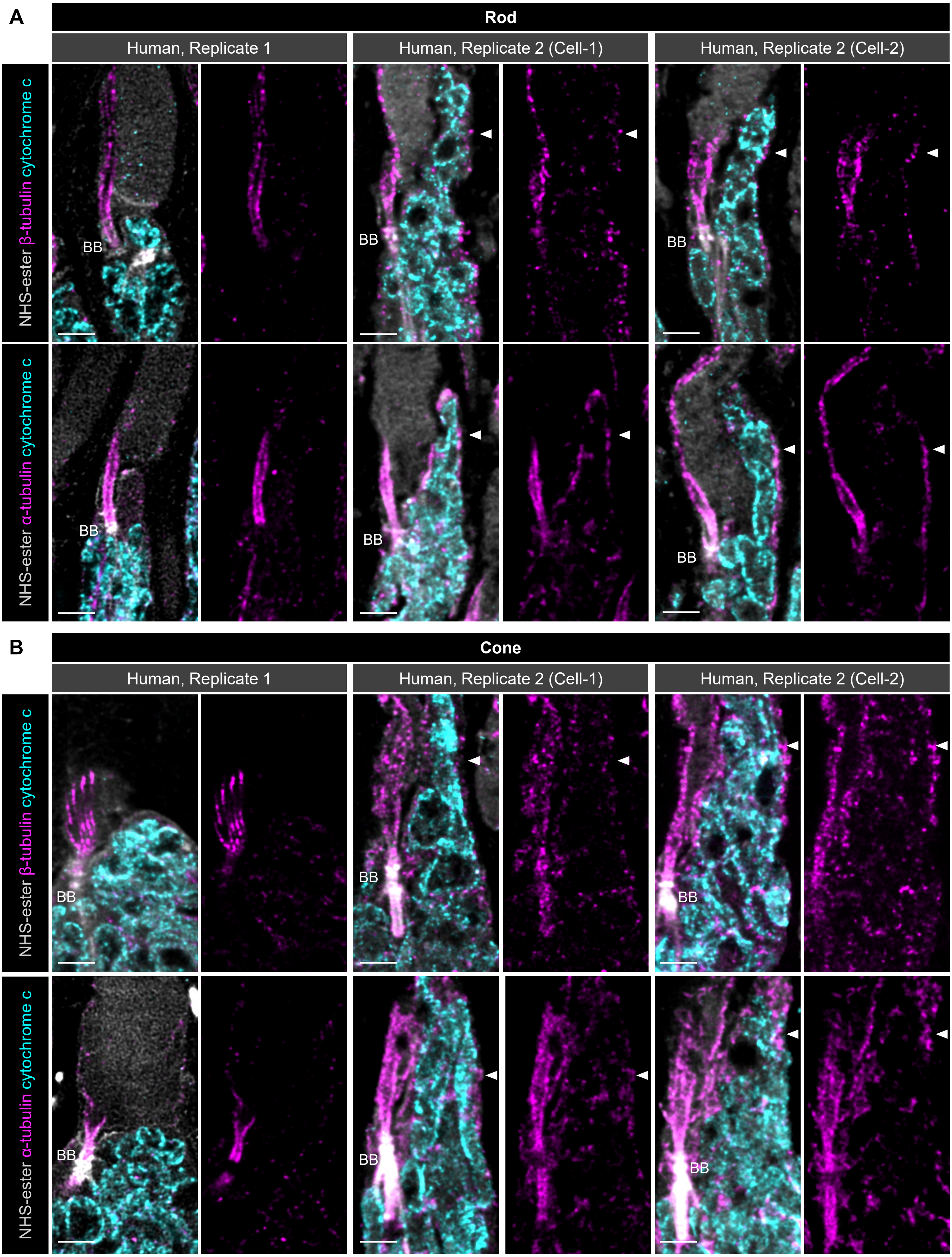
A
Rod
Human, Replicate 1
Human, Replicate 2 (Cell-1)
Human, Replicate 2 (Cell-2)
NHS-ester β-tubulin cytochrome c
NHS-ester α-tubulin cytochrome c
BB
B
Cone
Human, Replicate 1
Human, Replicate 2 (Cell-1)
Human, Replicate 2 (Cell-2)
NHS-ester β-tubulin cytochrome c
NHS-ester α-tubulin cytochrome c
BB

**Fig. S8. Tubulin organization in human accessory inner segment-like structures.**

**(A, B)** Representative single optical section images of human rod (A) and cone (B) photoreceptors showing the distribution of IS mitochondria visualized by cytochrome c immunolabeling (cyan) and the microtubule cytoskeleton visualized by β-tubulin (upper panels) or α-tubulin (lower panels) immunolabeling (magenta). In human replicate 1, aIS-like structures and associated tubulin-positive extensions were not observed in either rods or cones. In human replicate 2, several rods and cones showed mitochondria-containing aIS-like structures extending alongside the OS. Both β-tubulin- and α-tubulin-positive signals were detected within these aIS-like regions, but the signals appeared variable and uneven, resembling the broader IS microtubule network rather than a scaffold unique to the aIS-like structure. One representative cell is shown for each staining condition and rods and cones in human replicate 1, and two representative cells are shown for each staining condition and rods and cones in human replicate 2. All images are shown as single optical sections. Scale bars, 1 μm, corrected for the expansion factor. All microscopy images in this supplementary figure are U-ExM images.

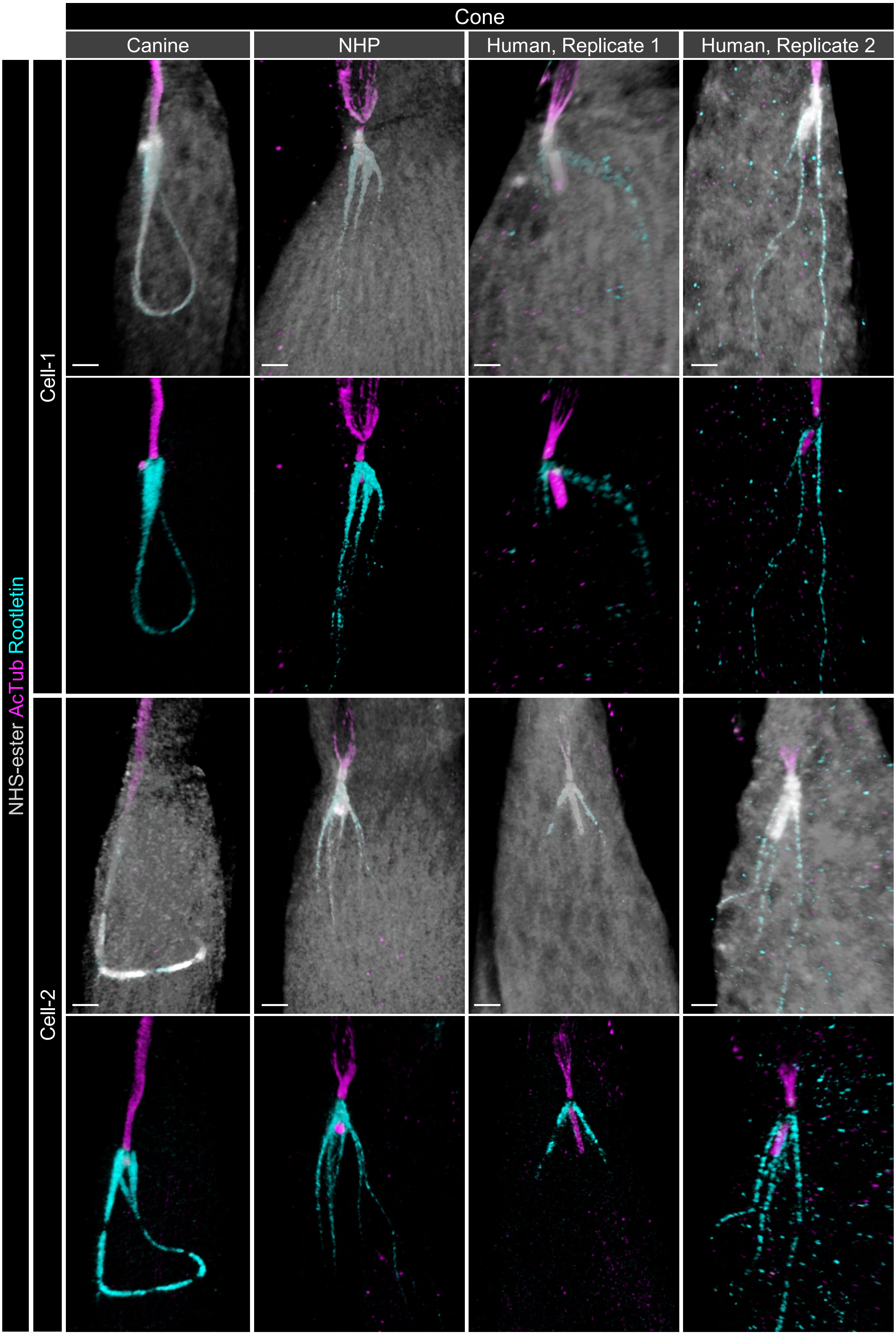
Cone
Canine
NHP
Human, Replicate 1
Human, Replicate 2
Cell-1
Cell-2
NHS-ester AcTub Rootletin

**Fig. S9. Additional examples of cone ciliary rootlet organization across species.**

Representative U-ExM images of cone photoreceptors immunolabeled for rootletin, showing additional examples not included in Fig. 8B. Two cone photoreceptors are shown for each group: canine, NHP, human replicate 1, and human replicate 2. In canine cones, the terminal portions of the ciliary rootlet form a continuous arc-like configuration. In NHP and human cones, the rootlet termini are separated into thin fibers extending toward the basal IS region. Scale bars, 1 µm, corrected for the expansion factor. All microscopy images in this figure are U-ExM images.

**Supplementary Table S1: List of archival retinas used in this study.**

| Species | ID | Sex | Age | Eye | Postmortem interval / Fixation conditions | Ref. |
|---|---|---|---|---|---|---|
| Canine | RC702 | M | 24 weeks | OS | < 5 mins / 4% FA 3 hrs, 2% FA 24 hrs | Takahashi et al., 2026 |
| | RC703 | M | 24 weeks | OS | | |
| | RC706 | F | 24 weeks | OS | | |
| NHP (cynomolgus macaques) | 21946 | F | 4 years | OD | < 20 mins / 4% FA 3 hrs, 2% FA 24 hrs | Jacobson et al., 2007 |
| | 21961 | F | 4 years | OS | | |
| Human | 125546 ODWE (replicate 1) | M | 83 years | OS, OD | ~ 3 hrs / 4% FA 7 days | NA |
| | 123991 ODWE (replicate 2) | F | 84 years | OS | ~ 12 hrs / 4% FA 24 hrs | NA |

F, female; FA, formaldehyde; M, male; NHP, non-human primate; OD, right eye; OS, left eye.

**Supplementary Table S2. Reagents and materials used for tissue processing and U-ExM.**

| Product | Supplier | Cat# |
|---|---|---|
| **Tissue processing and storage** | | |
| 20% paraformaldehyde solution | Electron microscopy sciences | 15713-S |
| Sucrose | Fiser chemical | S2-500 |
| Tissue-Tek O.C.T. Compound | Sakura finetek | 4583 |
| Tissue Embedding Disposable Molds | EBSciences | H1513 |
| **U-ExM** | | |
| 14-mm microwell/35-mm petri dish | MatTek | P35G-1.5-14-C |
| 12 mm Circular Cover Glasses | Fisher Scientific | 12541001 |
| Acrylamide (AA) | Sigma-Aldrich | A4058 |
| Ammonium Persulfate (APS) | Bio-Rad | 1610700 |
| Formaldehyde solution | Sigma-Aldrich | F8775 |
| ImmEdge Pen | Vector laboratories | H-4000 |
| Nuclease-Free Water | Invitrogen | AM9937 |
| N, N′-methylenbisacrylamide (BIS) | Sigma-Aldrich | M1533 |
| Phosphate Buffered Saline (PBS), 10x | Bio-Rad | 161-0780 |
| Poly-D-Lysine | Gibco | A3890401 |
| Sodium Acrylate (SA) | Sigma-Aldrich | 408220 |
| Sodium Chloride (NaCl) | Fisher Chemical | S271-3 |
| Sodium Dodecyl Sulfate (SDS) | Fisher Chemical | BP166-500 |
| Tetramethylethylenediamine (TEMED) | Bio-Rad | 161-0800 |
| Tris Base | Fisher Chemical | BP152-5 |

**Supplementary Table S3: Antibodies and regents used for immunolabeling.**

**Primary antibodies**

| Antibody | Host Organism | Clonality / Isotype | Cat# or Ref. | Dilution | RRID |
|---|---|---|---|---|---|
| Acetylated α-tubulin | Mouse | Monoclonal $IgG_{2b}$ | T7451 | 1/1000 | AB_609894 |
| Acetylated α-tubulin | Rabbit | Monoclonal IgG | ab179484 | 1/1000 | AB_2890906 |
| α-tubulin | Rabbit | Monoclonal IgG | ab18251 | 1/500 | AB_2210057 |
| β-actin | Rabbit | Polyclonal IgG | ab8227 | 1/200 | AB_2305186 |
| Blue opsin | Rabbit | Polyclonal IgG | AB5407 | 1/500 | AB_177457 |
| β-tubulin | Mouse | Monoclonal $IgG_1$ | T4026 | 1/500 | AB_477577 |
| CEP164 | Rabbit | Polyclonal IgG | 22227-1-AP | 1/200 | AB_2651175 |
| CEP290 | Rabbit | Polyclonal IgG | 22490-1-AP | 1/200 | AB_10973679 |
| cytochrome c | Sheep | Polyclonal IgG | C9616 | 1:200 | AB_532232 |
| Glutamylation (GT335) | Mouse | Monoclonal $IgG_1$ | AG-20B-0020 | 1/1000 | AB_2490210 |
| LCA5 (lebercilin) | Rabbit | Polyclonal IgG | 19333-1-AP | 1/200 | AB_2878576 |
| PCDH15 | Sheep | Polyclonal IgG | AF6729 | 1/300 | AB_10892338 |
| POC5 | Rabbit | Polyclonal IgG | A303-341A | 1/200 | AB_10971172 |
| Red/Green opsin | Rabbit | Polyclonal IgG | AB5405 | 1:300 | AB_177456 |
| Rhodopsin | Rabbit | Polyclonal IgG | AB9279 | 1/1000 | AB_11210489 |
| Rootletin | Human | Monoclonal IgG | HCA009 | 1/200 | AB_2085504 |
| Rootletin | Mouse | Monoclonal $IgG_1$ | sc-374056 | 1/200 | AB_10918081 |
| RP1 | Chicken | Polyclonal IgY | Custom (Liu et al., 2002) | 1/300 | NA |
| SPATA7 | Rabbit | Polyclonal IgG | 12020-1-AP | 1/200 | AB_2195380 |
| Whirlin | Rabbit | Polyclonal IgG | 25881-1-AP | 1/200 | AB_2880280 |

**Secondary antibodies**

| Target | Host Organism | Clonality / Isotype | Fluorescent | Cat# | Dilution | RRID |
|---|---|---|---|---|---|---|
| Chicken IgY (H+L) | Goat | Polyclonal IgG | Alexa Fluor 488 | A11039 | 1:1000 | AB_2534096 |
| Mouse $IgG_1$ | Goat | Polyclonal IgG | Alexa Fluor 488 | A21121 | 1:1000 | AB_2535764 |
| Mouse $IgG_{2b}$ | Goat | Polyclonal IgG | Alexa Fluor 488 | A21141 | 1:1000 | AB_2535778 |
| Sheep IgG (H+L) | Donkey | Polyclonal IgG | Alexa Fluor 488 | A11015 | 1:1000 | AB_2534082 |
| Human IgG (H+L) | Goat | Polyclonal IgG | Alexa Fluor 568 | A21090 | 1:1000 | AB_2535746 |
| Mouse $IgG_1$ | Goat | Polyclonal IgG | Alexa Fluor 568 | A21124 | 1:1000 | AB_2535766 |
| Mouse IgG (H+L) | Donkey | Polyclonal IgG | Alexa Fluor 568 | A10037 | 1:1000 | AB_11180865 |
| Rabbit IgG (H+L) | Goat | Polyclonal IgG | Alexa Fluor 568 | A11036 | 1:1000 | AB_10563566 |
| Rabbit IgG (H+L) | Donkey | Polyclonal IgG | Alexa Fluor 568 | A10042 | 1:1000 | AB_2534017 |

**Fluorescent conjugates**

| Product | Fluorescent | Supplier | Cat# | Stock solution | Dilution |
|---|---|---|---|---|---|
| Hoechst 33342 | Hoechst 33342 | Invitrogen | 62249 | 20 mM | 1:5000 |
| NHS-ester | Atto 647N | Sigma-Aldrich | 18373 | 2 mg/mL | 10 µg/mL |
| PNA | Alexa Fluor 568 | Invitrogen | L32458 | 1 mg/mL | 10 µg/mL |
| WGA | Alexa Fluor 488 | Invitrogen | W11261 | 1 mg/mL | 10 µg/mL |

RRID, research resource identifiers.
